# The Apolipoprotein E N-Terminal Bundle Rendered Membrane-Compatible by Reverse-QTY Conversion Without Loss of Fold

**DOI:** 10.64898/2026.09.25.754306

**Authors:** Taner Karagöl, Alper Karagöl

## Abstract

Apolipoprotein E (ApoE) performs most of its pathologically relevant biology at lipid interfaces, yet design efforts target its receptor binding, interdomain geometry or abundance rather than the lipid affinity of its N-terminal bundle. We asked whether the reverse-QTY (rQTY) code can make that bundle membrane-compatible without disturbing its fold. Within mature residues 11–167 we converted every glutamine, threonine and tyrosine lying inside an α-helix (Q→L, T→V, Y→F) and left those in turns and loops unchanged: 22 substitutions over 14.0% of the segment, sparing the receptor recognition region, moving the grand average of hydropathicity from −0.757 to +0.123. Across five AlphaFold3 models per sequence the variant bundle superimposes on the native at 0.81 Å mean Cα root-mean-square deviation with indistinguishable helix content, while apolar solvent-accessible surface rises 59% at constant total surface and burial. Hydrophobic moment increases in five of six helical segments, so amphipathicity is preserved. In all-atom molecular dynamics the native segment holds its fold in water over 100 ns; the variant stays folded and essentially fully helical in a six-component neuronal bilayer (50 ns) and a highly mobile membrane mimetic (90 ns), thinning the bilayer locally by 10.9 Å. The first lipid shell is only mildly biased in composition, and the two membrane models disagree. Each system was run once with the protein embedded at the build stage, so quantifying affinity will require matched native trajectories and partitioning free-energy calculations. rQTY thus acts as a geometry-preserving, residue-resolved control on the surface chemistry of a soluble helical bundle.

## Introduction

Apolipoprotein E (ApoE) is the principal lipid transport protein of the central nervous system. It is synthesised predominantly by astrocytes, and to a lesser extent by microglia and stressed neurons, and is secreted as the scaffolding component of discoidal high-density-lipoprotein-like particles whose assembly requires ATP-binding cassette transporter A1 (ABCA1)-mediated lipidation [1, 2]. Through these particles ApoE distributes cholesterol and phospholipids to neurons [3] for membrane biogenesis, synaptogenesis and axonal repair, and returns lipid cargo to cells through receptors of the low-density lipoprotein receptor (LDLR) family [1, 3]. The human *APOE* gene is polymorphic, and its three common alleles (ε2, ε3, ε4) encode isoforms that differ at only two positions, residues 112 and 158 of the mature protein [1, 4]. Despite this minimal chemical difference, the ε4 allele is the strongest common genetic risk factor for late-onset Alzheimer’s disease, acting in an allele dose-dependent manner on both lifetime risk and age at onset [5], with the magnitude of the association further modified by age, sex and ancestry [6]. The ε2 allele, conversely, is protective [1, 4].

The mechanisms that connect ApoE4 to neurodegeneration have proven considerably broader than amyloid deposition alone. Human ApoE isoforms differentially regulate the clearance of amyloid-β from the brain interstitial fluid [7], but ApoE4 also markedly exacerbates tau-mediated neurodegeneration in the absence of amyloid pathology [8], and *APOE4/4* microglia from patients accumulate damaging lipid droplets and adopt a dysfunctional, pro-inflammatory lipid-handling state [9]. Cerebrovascular integrity, myelination and glial immune signalling are likewise affected in an isoform-dependent manner [1, 4]. What these otherwise disparate mechanisms share is that each is executed at a lipid interface: the loading and unloading of lipoprotein particles, the extraction of lipid from membranes, the delivery of lipid to receptors, and the disposal of lipid within glia. The physical interaction between ApoE and lipid surfaces is therefore not a peripheral biophysical detail of the protein, but the substrate on which most of its pathologically relevant biology operates.

This interaction is governed by a two-domain architecture. The mature 299-residue protein comprises an N-terminal domain (residues 1–191) and a C-terminal domain (residues 192–299, whose high-affinity lipid-binding determinants lie in the region around residues 216–299) joined by a protease-sensitive, conformationally flexible hinge [10, 11]. The N-terminal domain folds into an elongated, up-and-down four-helix bundle that buries its apolar faces in the bundle core and exposes basic surface patches, the largest of which carries the LDLR recognition determinants around residues 136–150 [10, 11]. Its helices belong to the amphipathic-helix family shared by the exchangeable apolipoproteins, in which apolar and charged residues segregate onto opposite faces of the helix cylinder and the resulting facial asymmetry, rather than bulk hydrophobicity, sets how the helix docks at a lipid surface [12]. The C-terminal domain also mediates self-association [11], and NMR analysis of full-length monomeric ApoE3 shows that the two domains pack against one another rather than behaving as independent units [13]. Productive lipid association requires the N-terminal bundle to open, so that helices which are mutually packed in the lipid-free state can present their apolar faces to the acyl chain region [11]. Cryo-electron microscopy of ApoE secreted by astrocytes shows that on discoidal particles the lipidated protein is arranged as antiparallel dimers, so the conformation that engages lipid in a physiological particle is substantially rearranged relative to the closed, lipid-free bundle [14]. Isoform-specific behaviour is superimposed on this mechanism: in ApoE4 an Arg61–Glu255 domain interaction reorients the C-terminal domain against the N-terminal bundle and biases particle preference and lipidation state [15]. The conformational accessibility of the N-terminal helices to a bilayer is, in other words, a functional switch rather than an incidental property.

Attempts to make ApoE pharmacologically tractable have so far approached this switch indirectly. ApoE-mimetic peptides derived from the receptor-binding region, such as COG1410, a modified 12-mer spanning residues 138–149, reproduce receptor engagement and anti-inflammatory signalling in injury models [16]. Building on the observation that ApoE3 and ApoE4 differ conformationally [17], small-molecule structure correctors were developed to disrupt the ApoE4 domain interaction and drive ApoE4 towards an ApoE3-like conformation [18]. Antibody, antisense and expression-lowering strategies act instead on the abundance or extracellular fate of the protein [4]. Each of these modulates receptor binding, interdomain geometry, or protein level. None treats the intrinsic affinity of the N-terminal helices for the bilayer as a designable parameter, even though that affinity determines whether and how the bundle opens. A method for retuning membrane affinity in a controlled, residue-resolved manner would provide a handle orthogonal to the existing approaches, and would allow the contribution of the lipid interface itself to be tested rather than inferred.

Systematic engineering of helix composition offers such a method. The QTY code replaces the hydrophobic residues leucine, isoleucine, valine and phenylalanine with glutamine, threonine and tyrosine, converting hydrophobic α-helices into hydrophilic ones while preserving helical geometry and the overall fold, and has been used to render integral membrane receptors and transporters water-soluble without detergents, with ligand binding and catalytic activity retained in the converted forms [19, 20, 21]. We have previously applied this design to neuronal membrane proteins, including the serotonin, dopamine and norepinephrine transporters, the glutamate transporters and their truncated isoforms [22], and eight synaptic vesicle proteins, and to bacterial integral membrane enzymes, where QTY variants predicted with AlphaFold2 [23] and AlphaFold3 [24] superimpose on their native counterparts at low root-mean-square deviation (RMSD) and retain native-like conformational behaviour in molecular dynamics simulations [25, 26, 27, 28, 29]. The same pairing of the code with AlphaFold3 has since been applied to the human aquaporin family, where analogues carrying substitutions at 43 to 49% of their transmembrane positions superimpose on the native structures below 0.6 Å [30]. Deep-learning design pipelines reach the same objective by a different route, inverting AlphaFold2 and redesigning sequences with ProteinMPNN to produce soluble analogues of claudins, rhomboid proteases and G-protein-coupled receptors that were validated crystallographically and functionally [31]. A fixed substitution rule is the less general of the two, but it is interpretable position by position, and the same substitution set transfers unchanged between sequence variants of one target. The inverse operation, the reverse-QTY (rQTY) code, applies the same pairings in the opposite direction (Q→L, T→V, Y→F) to increase hydrophobicity, and has been used to convert human serum albumin into self-assembling amphiphilic nanoparticles for anti-tumour drug delivery [32]. These exchanges are not synthetic curiosities. The L↔Q, I↔T and F↔Y pairs each correspond to a single nucleotide change at the second codon position, and we have catalogued these substitutions, together with the T↔V exchange, as natural variants in human genomic databases and across homologous sequences [26, 33], and have shown that the divergence between membrane-associated and soluble forms of a single receptor family leaves a detectable evolutionary signature in residue composition [34]. To date, however, rQTY has been used to construct amphiphiles *de novo*, and has not been applied to bias a pre-existing, functionally relevant lipid interaction within a soluble helix bundle.

Here we report a domain-restricted rQTY design of the ApoE N-terminal segment. Substitutions were confined to residues 11–167 of the mature protein, a segment that contains the complete four-helix bundle, and were applied to every glutamine, threonine and tyrosine residue located within an α-helix of that segment, leaving all such residues in turns, bends and connecting loops unchanged. This yielded 22 substitutions (14 Q→L, 4 T→V, 4 Y→F), corresponding to 14.0% of the segment, and shifted its grand average of hydropathicity (GRAVY) from −0.757 to +0.123 on the Kyte–Doolittle scale (Δ = 0.880) [35]. Because the design is restricted to the N-terminal domain, the C-terminal lipid-binding domain and the hinge remain unaltered, and the contiguous stretch containing the LDLR recognition region (residues 136–150) receives no substitution. We characterised the design at two levels. Sequence and structure were compared directly between the native and rQTY forms, using hydropathy and hydrophobic-moment profiling together with AlphaFold3 prediction of five models for each sequence [24], which together tested whether the conversion perturbs the fold. Conformational behaviour was then examined by all-atom molecular dynamics. The native segment was simulated in bulk water, which tested whether the isolated bundle holds its fold without the C-terminal domain, and the rQTY segment in two membrane environments: an explicit phospholipid bilayer of neuronal composition, and a highly mobile membrane mimetic (HMMM) with matched headgroup composition and a fluid acyl region [36]. Both membrane systems were constructed with the bundle already embedded, so they ask whether a helical bundle whose surfaces have been made hydrophobic can occupy a lipid environment without unfolding, and what it does to the membrane when it does.

## Results and Discussion

### Helix-Restricted Design of the rQTY Variant

The design target was the segment spanning residues 11–167 of mature ApoE (157 residues), which contains the complete four-helix bundle of the N-terminal domain and terminates before the hinge. The reverse-QTY code was applied under a single secondary-structure rule, stated in full in Methods: convert every glutamine, threonine and tyrosine lying within an α-helix (Q→L, T→V, Y→F) and leave every such residue in a turn, bend or connecting loop unchanged. Because no criterion of burial, conservation or helical face enters, the rule is fully specified by the DSSP assignment and regenerates the variant sequence deterministically from the native one.

Applying it yielded 22 substitutions: 14 Q→L, 4 T→V and 4 Y→F, at mature positions 16, 17, 18, 36, 41, 46, 48, 55, 57, 58, 67, 74, 81, 89, 98, 101, 117, 118, 123, 156, 162 and 163 (Table 1, Fig. 1a). This corresponds to 14.0% of the segment, leaving 86.0% sequence identity to the native protein. All 22 substituted positions fall inside DSSP-assigned helices, and all six glutamine, threonine and tyrosine residues that were not substituted (Gln21, Gln24, Thr42, Thr83, Gln128, Thr130) lie outside them, at helix caps or in the loops that connect the bundle helices (Supplementary Table S1). The substitution rate is markedly lower than in our previous forward-QTY designs of integral membrane proteins, where 45–55% of transmembrane residues are typically replaced [26, 27, 28]; the difference reflects the starting material, since a soluble bundle presents far fewer convertible positions per helix than a transmembrane helix presents in the opposite direction.

**Figure 1.**
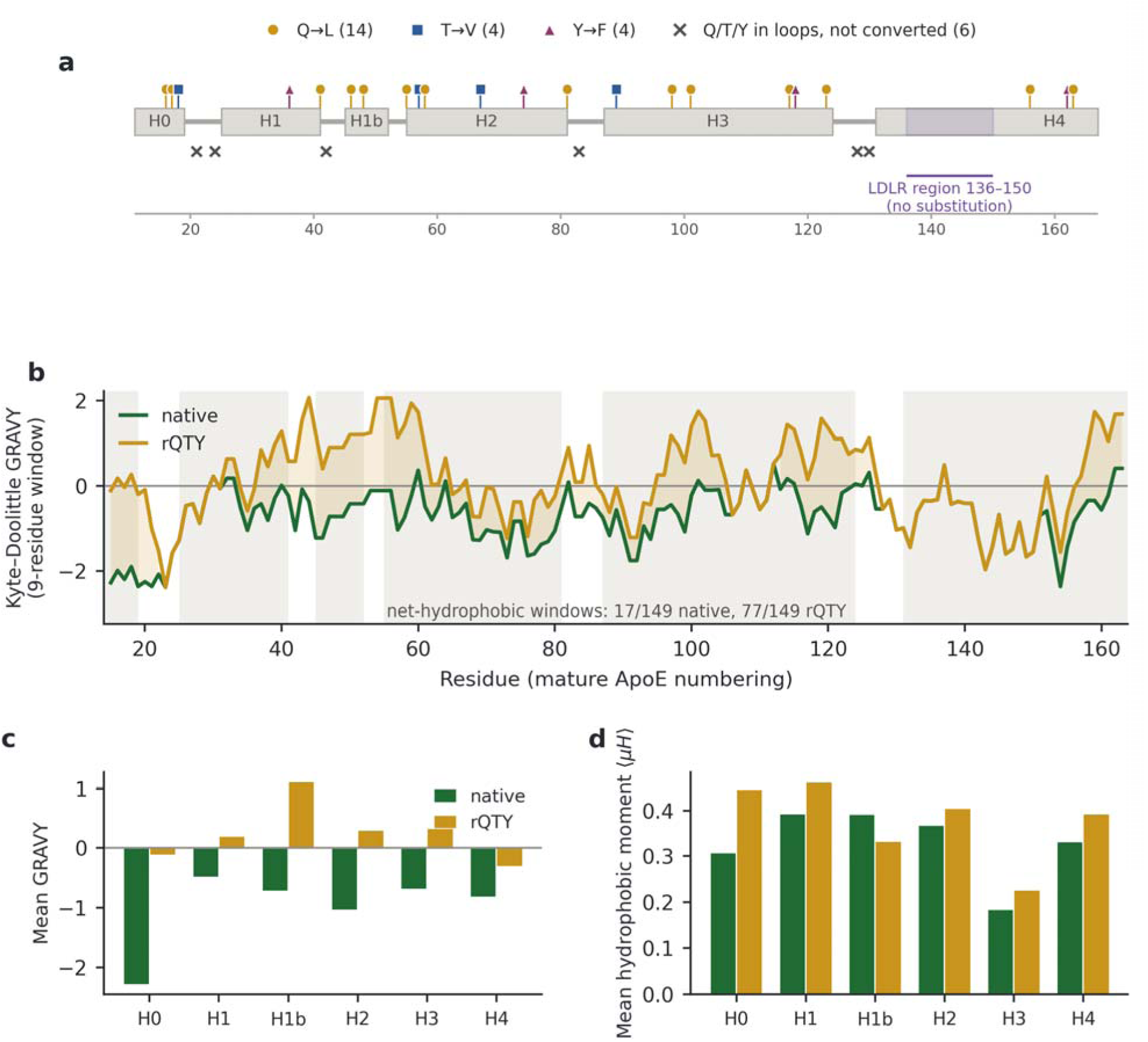
Design of the reverse-QTY variant of the ApoE N-terminal segment. (a) Position of the 22 rQTY substitutions along mature residues 11–167. Grey boxes are the α-helical segments assigned by DSSP; lollipops above the track mark converted positions, coloured and shaped by substitution type (Q→L gold circles, T→V slate squares, Y→F plum triangles); crosses below mark the six glutamine, threonine and tyrosine residues that lie outside helices and were therefore left unchanged. The purple block marks the LDLR recognition region (136–150), which receives no substitution. (b) Kyte–Doo ittle hydropathy in a nine-residue sliding window for the native (green) and rQTY (gold) sequences; grey shading marks the helical segments and the gold fill the local increase in hydropathy. (c) Mean Kyte– Doolittle GRAVY per helical segment. (d) Mean hydrophobic moment μH per helical segment, computed on the Eisenberg consensus scale at 100° per residue. Segment boundaries in (c) and (d) are those listed in Table 2.

**Table 1.** The 22 reverse-QTY substitutions in the ApoE N-terminal segment. Positions use mature ApoE numbering. DSSP assignment is taken from the full-length model used for the design; RSA is relative solvent accessibility averaged over the five AlphaFold3 models of each sequence. The complete list of all glutamine, threonine and tyrosine residues in the segment, including the six that were not converted, is given in Supplementary Table S1.

| Position | Native | rQTY | Helix | DSSP | RSA nat (%) | RSA rQTY (%) |
| --- | --- | --- | --- | --- | --- | --- |
| 16 | Q16 | L16 | H0 | H | 22.3 | 34.3 |
| 17 | Q17 | L17 | H0 | H | 18.4 | 32.7 |
| 18 | T18 | V18 | H0 | H | 49.8 | 48.8 |
| 36 | Y36 | F36 | H1 | H | 14.6 | 16.7 |
| 41 | Q41 | L41 | H1 | H | 11.3 | 19.8 |
| 46 | Q46 | L46 | H1b | H | 46.0 | 45.0 |
| 48 | Q48 | L48 | H1b | H | 20.6 | 18.2 |
| 55 | Q55 | L55 | H2 | H | 42.9 | 47.5 |
| 57 | T57 | V57 | H2 | H | 12.8 | 10.7 |
| 58 | Q58 | L58 | H2 | H | 51.5 | 56.1 |
| 67 | T67 | V67 | H2 | H | 0.0 | 0.1 |
| 74 | Y74 | F74 | H2 | H | 10.3 | 7.9 |
| 81 | Q81 | L81 | H2 | H | 55.8 | 33.5 |
| 89 | T89 | V89 | H3 | H | 31.0 | 34.0 |
| 98 | Q98 | L98 | H3 | H | 50.4 | 47.6 |
| 101 | Q101 | L101 | H3 | H | 10.8 | 5.4 |
| 117 | Q117 | L117 | H3 | H | 42.3 | 48.7 |
| 118 | Y118 | F118 | H3 | H | 1.9 | 2.2 |
| 123 | Q123 | L123 | H3 | H | 55.4 | 51.9 |
| 156 | Q156 | L156 | H4 | H | 18.2 | 25.9 |
| 162 | Y162 | F162 | H4 | H | 5.0 | 1.1 |
| 163 | Q163 | L163 | H4 | H | 26.3 | 25.6 |

**Table 2.** Hydropathy and amphipathicity by segment. Segments are the α-helices assigned by DSSP on the full-length model; residues in the connecting loops are excluded from the per-segment rows but included in the whole-segment row. GRAVY uses the Kyte–Doolittle scale; H and μH are the mean hydrophobicity and mean hydrophobic moment on the Eisenberg consensus scale, computed at 100° per residue.

| Segment | Residues | n | Subs | GRAVY<br>nat | GRAVY<br>rQTY | $\langle H \rangle$ nat | $\langle H \rangle$<br>rQTY | $\langle \mu H \rangle$ nat | $\langle \mu H \rangle$<br>rQTY |
| --- | --- | --- | --- | --- | --- | --- | --- | --- | --- |
| H0 | 11–19 | 9 | 3 | −2.278 | −0.111 | −0.591 | −0.041 | 0.306 | 0.445 |
| H1 | 25–41 | 17 | 2 | −0.482 | 0.188 | −0.049 | 0.118 | 0.392 | 0.462 |
| H1b | 45–52 | 8 | 2 | −0.713 | 1.113 | −0.090 | 0.388 | 0.390 | 0.331 |
| H2 | 55–81 | 27 | 6 | −1.033 | 0.293 | −0.263 | 0.067 | 0.366 | 0.404 |
| H3 | 87–124 | 38 | 6 | −0.682 | 0.324 | −0.249 | 0.007 | 0.183 | 0.225 |
| H4 | 131–167 | 37 | 3 | −0.814 | −0.308 | −0.452 | −0.324 | 0.330 | 0.392 |
| Whole | 11–167 | 157 | 22 | −0.757 | 0.123 | −0.231 | −0.009 | 0.094 | 0.109 |

Two features of the resulting variant are consequences of the rule rather than of explicit design, and both are favourable. First, the substituted positions are distributed unevenly along the sequence: there is no substitution anywhere between residues 124 and 155, so the contiguous stretch containing the LDLR recognition determinants, residues 136–150, is entirely native [10, 11]. Drawn on the fold, not one of the 22 side chains falls on that helix (Fig. 2a): the receptor-binding surface is preserved by construction, not by intervention. Second, because the conversion is confined to the N-terminal domain, the C-terminal high-affinity lipid-binding region and the interdomain hinge are untouched; in a full-length context they would remain wild-type, and only the bundle’s own interaction with lipid would be altered. These are exactly the boundary conditions required if the aim is to interrogate the N-terminal bundle as an independent membrane-binding module rather than to rebuild the protein.

**Figure 2.**
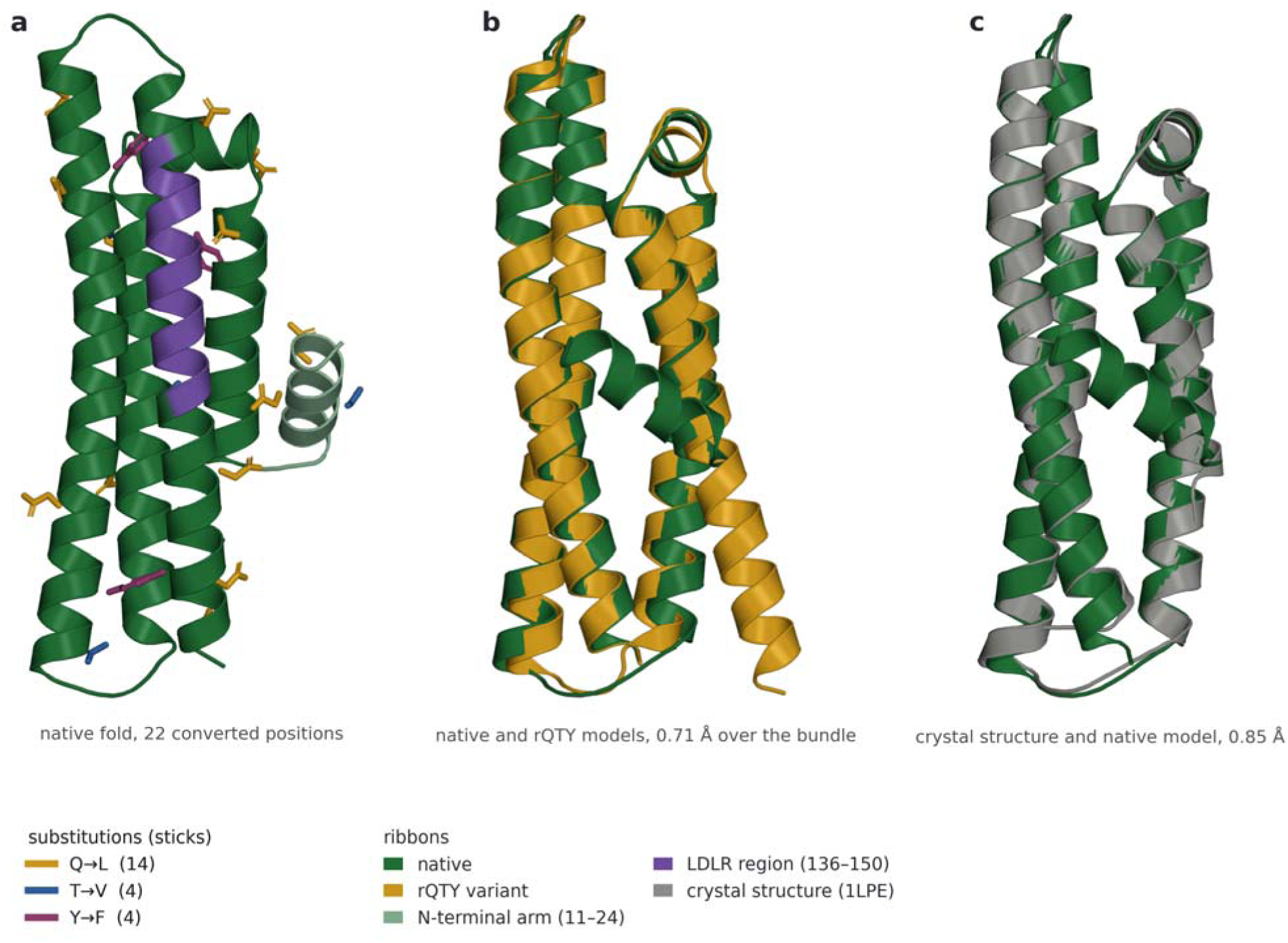
The designed fold. (a) The 22 converted positions on the top-ranked AlphaFold3 model of the native segment, drawn as side-chain sticks and coloured by substitution type as in Fig. 1a; the bundle is green, the N-terminal arm (residues 11–24) is the paler green and the LDLR recognition region (136– 150) is violet. No stick falls on the violet helix, which is the geometric statement of the rule sparing the receptor determinants. (b) The top-ranked native (green) and rQTY (gold) models superposed on the bundle alone, residues 25–167, at 0.71 Å over 143 Cα atoms; the arm is left free, because its placement is what differs between the two ensembles and not the packing of the bundle. (c) The crystal structu e of the wild-type ApoE3 N-terminal domain (1LPE, grey) superposed on the same native model over residues 25–166, at 0.85 Å over 142 Cα atoms, which is the agreement between the model the design was taken from and experiment. All three panels share one orientation and are ray traced orthoscopically.

The assignment the rule depends on is a property of one structure, so the substitution set was re-derived from two experimental structures of the same protein (Supplementary Tables S4 and S5). The model itself stands up to that comparison: superposed on the crystal structure of the wild-type ApoE3 N-terminal domain over residues 25–166, it agrees to 0.85 Å across 142 Cα atoms (Fig. 2c), so the fold the design was read from is the experimental fold. Helix interiors are unanimous: 23 of the 28 glutamine, threonine and tyrosine positions in the segment receive the same decision from every structure that resolves them. The crystal structure reproduces the boundaries of H1 (25–41) and H1b (45–52) exactly and agrees with the rule at 22 of the 24 positions it contains; the NMR ensemble agrees at 23 of 28. What differs is three helix ends, and at each of them the two experimental structures agree with each other rather than with the prediction: H2 ends at residue 78 rather than 81, H4 at 162 rather than continuing to the segment boundary, and H0 extends to 22 rather than stopping at 19. Three decisions follow from those three boundaries. Q81 and Q163 were converted on a helical assignment that neither experimental structure supports, and Q21 was left unconverted although it is helical in all 20 NMR conformers; two further positions, Q41 and T89, are helical in the crystal structure and in the design model but not in the majority of NMR conformers. The three substitutions in H0, at positions 16, 17 and 18, which the crystal structure does not resolve and which the design model places in its least confident region, are helical in all 20 conformers of the NMR structure. The design is therefore reproducible in helix interiors and model-dependent at helix termini, where two of the 22 conversions and one of the six retained positions would change had the assignment been taken from an experimental structure rather than from the prediction.

### Redistribution of Hydropathy and Amphipathicity

The conversion moves the grand average of hydropathicity (GRAVY, Kyte–Doolittle scale [35]) of the segment from −0.757 to +0.123, a shift of 0.880 that carries the construct across the neutral boundary from a net-hydrophilic to a net-hydrophobic sequence. The magnitude is comparable to the shifts we have reported in the opposite direction for forward-QTY variants of neuronal transporters [27, 28], indicating that the two operations are of similar chemical scale even though the substitution rate here is roughly a quarter as large.

The distribution of that change is more informative than its mean. In a nine-residue sliding window, the native segment contains 17 net-hydrophobic windows out of 149 (11%), spanning a range of −2.39 to +0.52; the rQTY segment contains 77 out of 149 (52%), spanning −2.39 to +2.07 (Fig. 1b). The lower bound is identical in the two sequences: the most hydrophilic stretches of ApoE, which contain no convertible residues, are unaffected. What the design does is create new hydrophobic windows rather than shift the whole profile, and the largest local increases (ΔGRAVY ≈ +2.2 to +2.4 over a nine-residue window) occur around residues 17–18, 44–45 and 57–58.

Resolving the same quantities by helix shows that the redistribution follows the architecture of the bundle (Table 2; mean GRAVY per segment in Fig. 1c). Mean hydrophobicity H on the Eisenberg consensus scale rises in every segment, but by very different amounts [37]: the short helix spanning residues 45–52 becomes the most hydrophobic of the six ( H −0.090 → +0.388; GRAVY −0.713 → +1.113 from only two substitutions in eight residues), although the largest shift on both scales is in H0 ( H −0.591 → −0.041; GRAVY −2.278 → −0.111), while helix H4 (131–167), which carries the receptor-binding region and receives only three substitutions, remains the most hydrophilic segment in the variant ( H −0.452 → −0.324; GRAVY −0.814 → −0.308). Critically, the mean hydrophobic moment μH does not collapse: it increases in five of six segments (for example H2, 0.366 → 0.404; H4, 0.330 → 0.392) and falls only modestly in the 45–52 helix (0.390 → 0.331) (Fig. 1d). The segment the conversion makes most hydrophobic is therefore also the one that loses facial asymmetry, which sets a practical limit on how far the rule can be pushed within a single short helix. Taken over the segment as a whole, the variant is nevertheless not a uniformly greasy helical bundle but a more hydrophobic one that retains, and in most segments slightly strengthens, the facial asymmetry characteristic of exchangeable apolipoprotein helices [12]. That distinction matters mechanistically, because it is amphipathicity rather than bulk hydrophobicity that governs how an apolipoprotein helix sits at a lipid– water interface [11, 12]. One caveat attaches to the moments themselves: those for H0 (nine residues) and H1b (eight residues) are computed over fewer residues than the eleven-residue window the method assumes, and are correspondingly less well determined.

### Fold Conservation in AlphaFold3 Predictions

Five AlphaFold3 [24] models were generated for each sequence. Global confidence is lower for the variant (predicted TM-score, pTM, 0.85 versus 0.79, equal in each case to the server ranking score; mean per-atom predicted local distance difference test score, pLDDT, 87.1 versus 80.9; mean predicted aligned error 5.19 Å versus 6.25 Å), and no model of either sequence was flagged for steric clash or predicted disorder. Confidence metrics do not measure stability, but a drop of this size would ordinarily prompt the question of whether the design has disrupted the fold. Resolving confidence by region shows where it comes from. Mean per-residue pLDDT over the bundle proper (residues 25–167) is high in both sequences, 90.7 for the native segment and 84.4 for the variant, whereas over the N-terminal arm (residues 11–24) it is 57.2 and 52.2 respectively. The arm is the low-confidence element in both sequences, and it is the region that drives the global statistics down (Fig. 3a).

**Figure 3.**
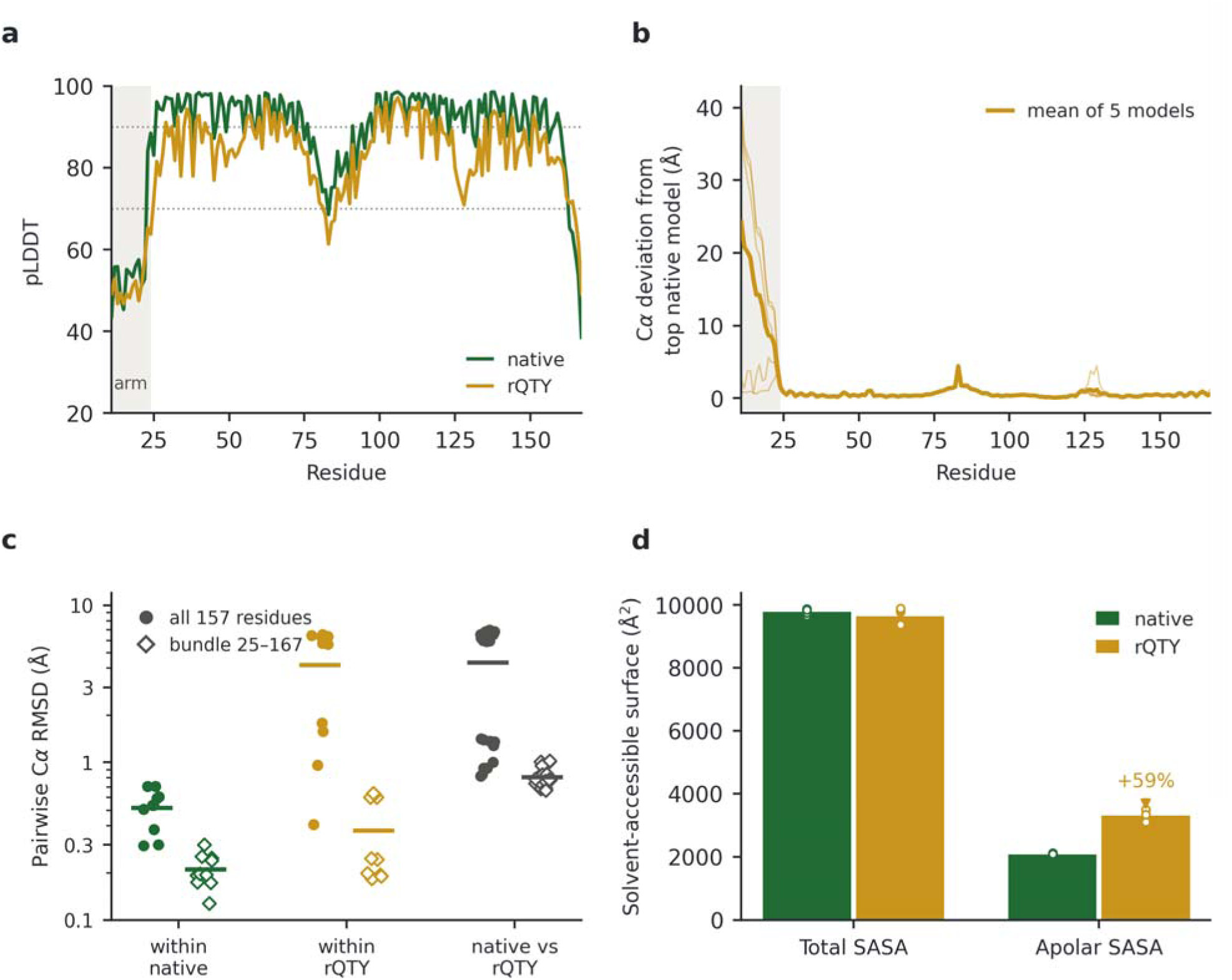
AlphaFold3 comparison of the native and rQTY segments, five models per sequence. (a) Perresidue pLDDT of the top-ranked model of each sequence; the shaded band is the N-terminal arm (residues 11–24), the low-confidence element in both sequences. Dashed lines mark pLDDT 70 and 90. (b) Per-residue Cα deviation of each of the five rQTY models from the top-ranked native model after superposition on the bundle (residues 25–167); the thick line is the mean of the five. (c) Pairwise Cα RMSD within the native ensemble (10 pairs), within the rQTY ensemble (10 pairs) and between the two ensembles (25 pairs), computed over all 157 residues (filled circles) and over the bundle alone (pen diamonds); horizontal bars are means. (d) Total and apolar solvent-accessible surface area (Shrake– Rupley); bars are ensemble means and points the five individual models.

Secondary structure is essentially identical. Helix content averaged over the five models is 84.6 ± 0.8% for the native segment and 85.6 ± 1.4% for the variant, and the helix boundaries coincide: in the segment models, 45–52, 55–81, 87–124 and 131–165 are predicted identically in both (the design boundaries in Table 2 come from the full-length model, in which H4 runs to residue 167). The single difference is at the N-terminus, where the native prediction resolves two helices (12–21 and 25–41) separated by a short break, while the variant predicts one continuous helix from 12 to 40. Three substitutions (Gln16→Leu, Gln17→Leu, Thr18→Val) lie within the arm itself and two more (Tyr36→Phe, Gln41→Leu) within H1, but the arm’s per-residue confidence (pLDDT 52–57) is too low for the difference in helix assignment to be interpreted.

Superposition quantifies how localised the structural consequences are. Over the full 157 residues, the native ensemble is tightly clustered (pairwise Cα root-mean-square deviation, RMSD, 0.52 Å, range 0.30–0.70 Å) while the variant ensemble splits into two groups (pairwise RMSD 4.18 Å, range 0.40– 6.52 Å): models R2 and R4 are native-like (0.82–1.41 Å from every native model), while R1, R3 and R5 form a second cluster 5.9–7.0 Å away (models are numbered R1–R5 throughout, as in Supplementary Fig. S1). Restricting the superposition to the bundle (residues 25–167) collapses the difference entirely. The native ensemble converges to 0.21 Å (0.13–0.30 Å), the variant to 0.37 Å (0.18–0.64 Å), and all 25 native-versus-variant pairs fall between 0.67 and 1.02 Å, with a mean of 0.81 Å (Fig. 3c; the complete pairwise matrices are given in Supplementary Fig. S1). The two top-ranked models superpose at 0.71 Å over the 143 bundle Cα atoms and are shown overlaid in Fig. 2b. Superposed on the bundle, the per-residue deviation between the two variant clusters averages 24.7 Å over residues 11–20 (maximum 38.9 Å), falls to 2.4 Å over residues 21–30, and does not exceed 2.5 Å anywhere beyond residue 30 (mean 0.23 Å); the corresponding deviation of each variant model from the top-ranked native model is shown in Fig. 3b. Distances between helix centroids are almost unchanged: H1–H2 is 11.9 Å in both ensembles, H3–H4 9.9 versus 9.8 Å, H1–H4 11.1 versus 11.7 Å, and the largest single change is at H2–H3 (12.4 versus 11.7 Å). The two variant clusters therefore differ in where the N-terminal arm lies, not in how the bundle is packed. The top-ranked variant model, R1, which was used to build both membrane systems, belongs to the cluster whose arm is displaced; the membrane trajectories therefore start from that arm placement rather than from the native-like one.

This is the central structural result and it is consistent with what the QTY code has repeatedly shown in the forward direction: replacing residues within an α-helix by their code partners changes the chemistry of the helical surface while leaving the helix, and the tertiary arrangement built from it, in place [19, 20, 26, 27, 28, 29]. The present data extend that observation to the reverse operation on a water-soluble helical bundle, and add a quantitative bound: every one of the 25 cross-ensemble bundle superpositions lies at or below 1.02 Å, mean 0.81 Å, after 22 substitutions. The one region that does respond, the low-confidence N-terminal arm, is precisely the region a prediction method should be least trusted on; the two AlphaFold3 clusters are better read as two placements of a mobile arm than as two folds.

### Apolar Surface at Constant Molecular Size

The conversion was intended to change what the bundle presents to solvent, and solvent-accessible surface area (SASA) computed by the Shrake–Rupley algorithm [38] shows that it does so with little change in molecular size. Total SASA differs by 1.3% between the two ensembles (9776 ± 82 Å² native versus 9646 ± 236 Å² variant), within the combined spread of the two ensembles, while apolar SASA rises from 2079 ± 32 Å² to 3313 ± 174 Å², an increase of 59%. The apolar fraction of the accessible surface therefore moves from 21.3 ± 0.2% to 34.3 ± 1.1% (Fig. 3d). The variant is not larger, more expanded, or more unfolded; the same surface has been repainted.

Relative solvent accessibility (RSA) at the 22 substituted positions is likewise almost unchanged by the substitution (27.2% in the native ensemble, 27.9% in the variant) [39], and remains slightly below the average for the rest of the chain (31.0% and 30.7%; Supplementary Fig. S2a). Two conclusions follow. First, the replacement side chains occupy the same degree of burial as the residues they replace. The Y↔F and T↔V pairs are both close to isosteric, the first removing a single hydroxyl oxygen from the aromatic ring and the second exchanging a hydroxyl for a methyl group at Cβ, whereas Q↔L exchanges a polar amide for an apolar isobutyl group of broadly similar bulk rather than of identical volume [20, 26]. Since 14 of the 22 substitutions are of the last kind, that burial is nonetheless unchanged is an empirical result of these models rather than a geometric necessity. Second, the design is not a surface-only operation. Ten of the 22 positions are solvent-exposed in the native fold (RSA > 25%) and four are strongly exposed (RSA > 50%), while others, among them Thr67, Tyr74, Tyr118, Tyr162 and Gln101, are largely buried in the bundle interior. The rule converts every helical Q, T and Y position regardless of burial: the exposed subset contributes the apolar surface gain, while the buried subset alters the packing chemistry of the core without changing its geometry.

### Conformational Stability of the Native Bundle in Water

The native segment was simulated in explicit water for 100 ns at 303.15 K. It is stable throughout. Cα RMSD from the starting structure plateaus at 1.94 ± 0.18 Å over the second half of the trajectory (1.88 ± 0.21 Å for the bundle alone), the radius of gyration is constant at 18.12 ± 0.09 Å (Fig. 4b), indistinguishable from the AlphaFold3 value of 18.05–18.14 Å, and mean Cα root-mean-square fluctuation (RMSF) is 1.01 Å, rising only to 1.25 Å over the N-terminal arm (Fig. 4, Table 3). The predicted bundle is thus metastable over 100 ns under an all-atom force field rather than dissipating as a prediction artefact, and the isolated 11–167 fragment does not require the C-terminal domain to hold its fold on this timescale. A single trajectory of this length establishes metastability, not the depth of the underlying free-energy minimum.

**Figure 4.**
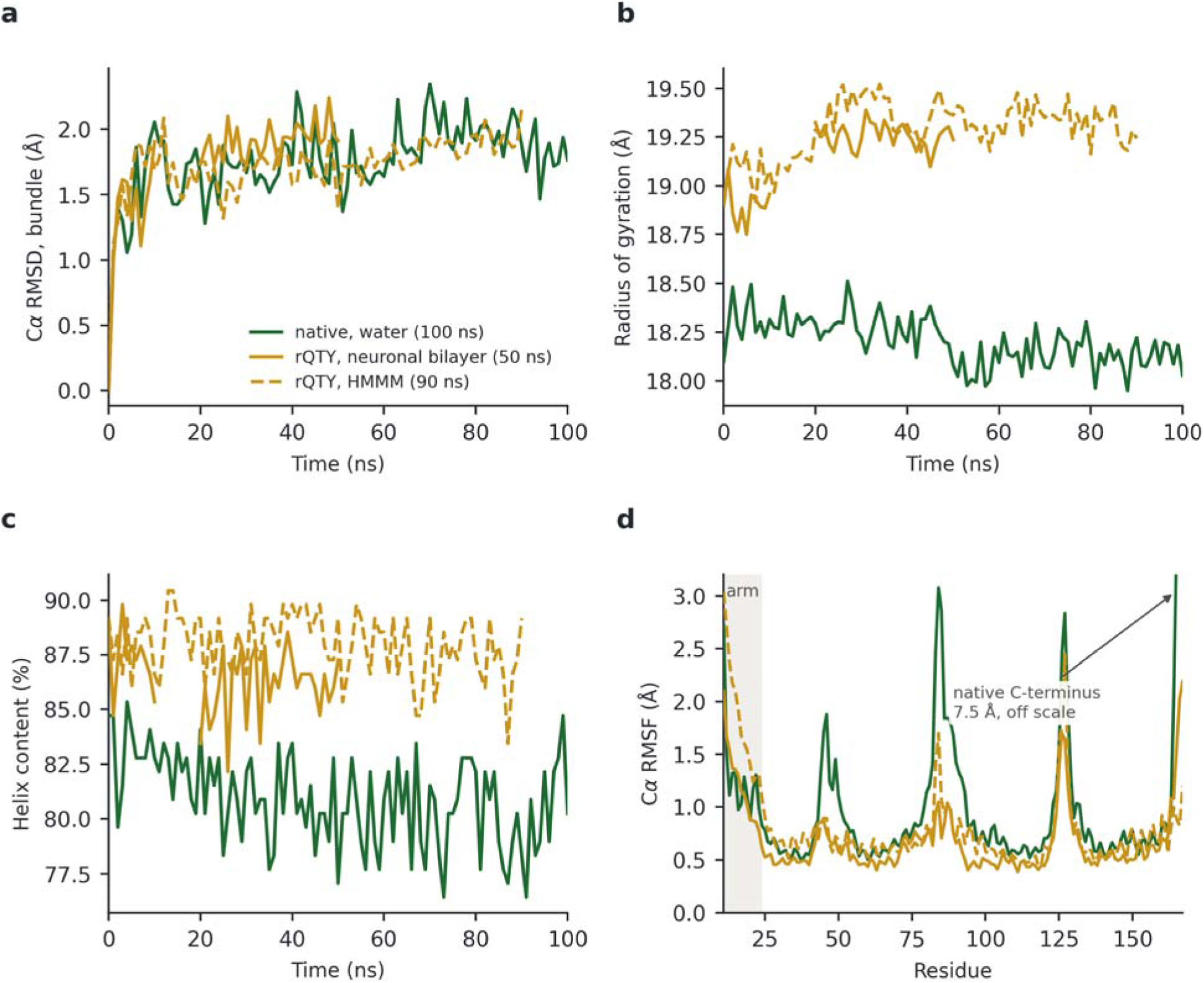
Conformational stability in all-atom molecular dynamics. (a) Cα RMSD of the bundle (residues 25–167) from the starting structure, (b) radius of gyration of the Cα trace and (c) DSSP helix content as functions of simulation time, for the native segment in water (green, 100 ns) and the r TY variant in the six-component neuronal bilayer (gold solid, 50 ns) and in the highly mobile membrane mimetic (gold dashed, 90 ns). The bilayer trace is broken between 10 and 20 ns, where the retained coordinates are unavailable (Supplementary Note S2). (d) Per-residue Cα RMSF over the second half of each trajectory after superposition on the bundle; the shaded band is the N-terminal arm. The vertical scale is truncated at 3.2 Å; the terminal residue of the native segment in water reaches 7.5 Å. Colours and line styles as in (a).

**Table 3.**
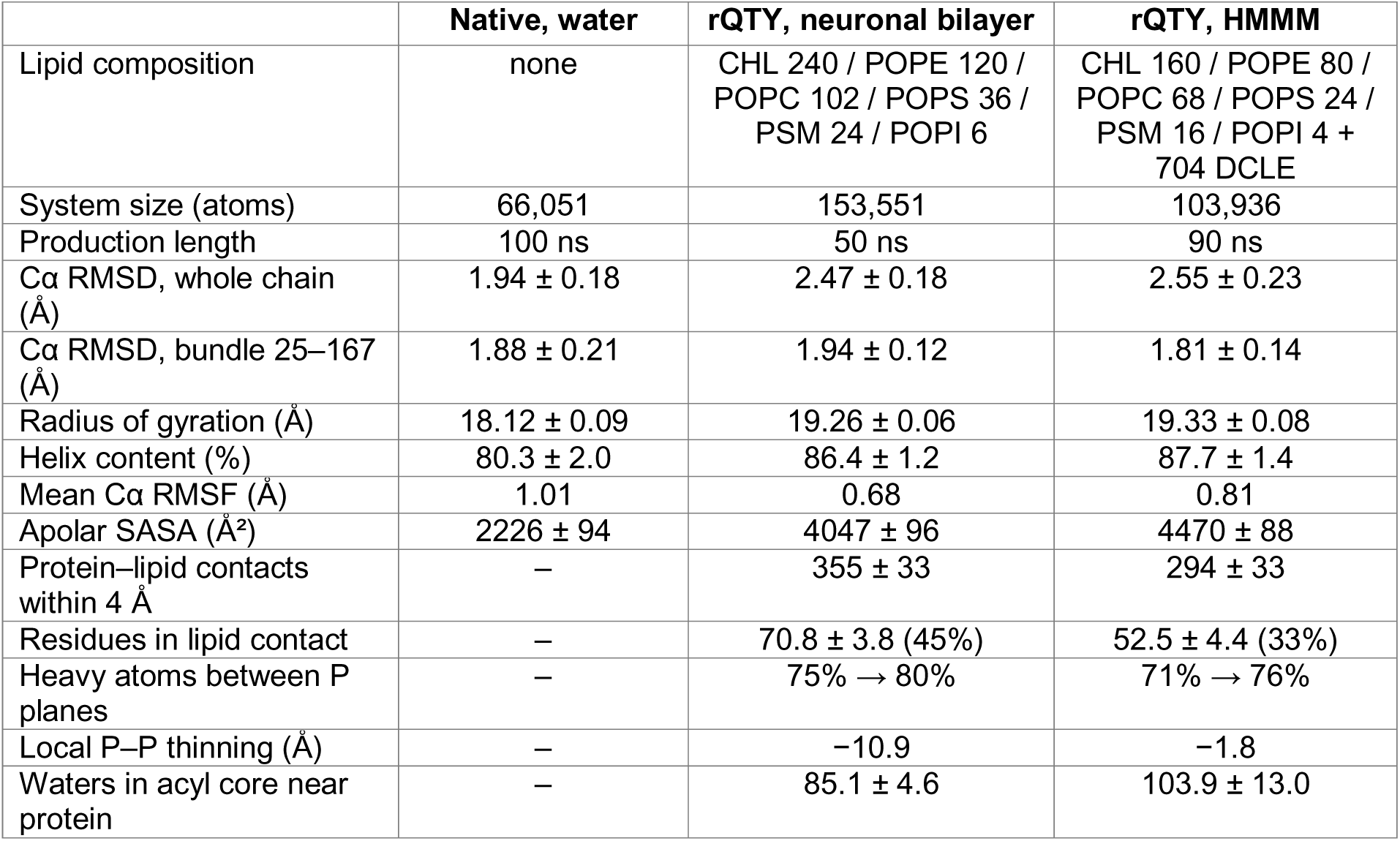
Molecular dynamics systems and equilibrated observables. All values are means ± standard deviation over the second half of each production run at 303.15 K. Dashes mark quantities that do not apply to the aqueous system. Local thinning is the difference between the 40–60 Å and 12–18 Å annuli of the same trajectory, computed from unrounded values. Each system was run once, so the tabulated deviations are the spread of a single equilibrated time series rather than standard errors over replicates. Bilayer values are computed over the 44 retained frames; see Supplementary Note S2. CHL, cholesterol; PSM, sphingomyelin; DCLE, 1,1-dichloroethane. Full system compositions are given in Supplementary Table S2.

|  | Native, water | rQTY, neuronal bilayer | rQTY, HMMM |
| --- | --- | --- | --- |
| Lipid composition | none | CHL 240 / POPE 120 / POPC 102 / POPS 36 / PSM 24 / POPI 6 | CHL 160 / POPE 80 / POPC 68 / POPS 24 / PSM 16 / POPI 4 + 704 DCLE |
| System size (atoms) | 66,051 | 153,551 | 103,936 |
| Production length | 100 ns | 50 ns | 90 ns |
| Cα RMSD, whole chain (Å) | 1.94 ± 0.18 | 2.47 ± 0.18 | 2.55 ± 0.23 |
| Cα RMSD, bundle 25–167 (Å) | 1.88 ± 0.21 | 1.94 ± 0.12 | 1.81 ± 0.14 |
| Radius of gyration (Å) | 18.12 ± 0.09 | 19.26 ± 0.06 | 19.33 ± 0.08 |
| Helix content (%) | 80.3 ± 2.0 | 86.4 ± 1.2 | 87.7 ± 1.4 |
| Mean Cα RMSF (Å) | 1.01 | 0.68 | 0.81 |
| Apolar SASA (Å<sup>2</sup>) | 2226 ± 94 | 4047 ± 96 | 4470 ± 88 |
| Protein–lipid contacts within 4 Å | – | 355 ± 33 | 294 ± 33 |
| Residues in lipid contact | – | 70.8 ± 3.8 (45%) | 52.5 ± 4.4 (33%) |
| Heavy atoms between P planes | – | 75% → 80% | 71% → 76% |
| Local P–P thinning (Å) | – | –10.9 | –1.8 |
| Waters in acyl core near protein | – | 85.1 ± 4.6 | 103.9 ± 13.0 |

Helix content declines modestly from 84.7% at the start to 80.3 ± 2.0% at equilibrium, and the loss is not evenly distributed. Resolved by segment, H1 (99.9%), H2 (97.6%) and H3 (95.7%) remain essentially fully helical, H4 is largely so (86.7%) and H0 falls to 77.8%, while the short 45–52 helix is helical for only 42.0% of the equilibrated trajectory. That element is a marginal helix in the native sequence, and its fraying accounts for the largest single part of the change. It is also, as noted above, the segment whose hydropathy the rQTY conversion alters most.

Apolar SASA over the equilibrated trajectory is 2226 ± 94 Å², 7.1% above the AlphaFold3 ensemble value of 2079 ± 32 Å² and within 1.5 combined standard deviations of it. That a static prediction and a 100 ns dynamic average agree this closely is a useful internal control for the native sequence, indicating that the surface-chemistry values reported in the preceding section are properties of the fold rather than of the modelling procedure. The equivalent control is not available for the variant, which was not simulated in water.

### The rQTY Bundle in a Neuronal-Composition Bilayer

The variant was placed in an explicit bilayer built to approximate a neuronal membrane (240 cholesterol, 120 POPE, 102 POPC, 36 POPS, 24 sphingomyelin and 6 POPI molecules, 528 lipids in total) and simulated for 50 ns. In this system the bundle is membrane-embedded rather than surface-associated: at the start of production 75% of protein heavy atoms lie between the two leaflet phosphate planes, and the protein spans from −32 Å to +35 Å relative to the bilayer midplane with its centre of mass at the midplane itself. The relevant question for the design is therefore not whether the variant partitions, but whether a four-helix bundle whose helical surfaces have been made hydrophobic can occupy that environment without unfolding.

It can, and it does so with the fold intact. Over 50 ns the fraction of protein heavy atoms between the phosphate planes rises from 75% to 80%, the centre of mass stays at the midplane (0.6 ± 0.6 Å over the second half), Cα RMSD plateaus at 2.47 ± 0.18 Å for the whole chain and 1.94 ± 0.12 Å for the bundle, and the radius of gyration is constant at 19.26 ± 0.06 Å (Fig. 4a, b, Table 3). Mean Cα RMSF is 0.68 Å, against 1.01 Å for the native protein in water, and the N-terminal arm, the most mobile element in every other measurement reported here, rises only to 1.16 Å, against 1.25 Å in the native run (Fig. 4d). Sequence and environment differ between the two runs, so the difference cannot be attributed to either alone. Helix content is 86.4 ± 1.2% (Fig. 4c), and every bundle helix is essentially fully formed: H1 98.8%, H2 98.5%, H3 96.8%, H4 91.2% and the 45–52 element 100.0%, with H0, the helical part of the N-terminal arm, at 77.8% (Supplementary Table S3). The marginal helix that frays in the native protein in water is fully helical in the variant in the membrane. Whether that reflects the raised GRAVY of the segment (−0.713 to +1.113), the change of environment, or both cannot be separated without a native trajectory in the same bilayer.

Engagement with lipid is extensive: 355 ± 33 protein–lipid heavy-atom contacts within 4 Å, involving 70.8 ± 3.8 of the 157 residues (45%, Fig. 5c); a snapshot of the embedded variant is shown in Fig. 5a. Per-residue depth in the membrane is given in Supplementary Fig. S2b. Contacts are distributed around the bundle rather than confined to one face (H1 53%, H2 56%, H3 55% and H4 38% of residues in contact), as expected for an embedded rather than a surface-bound orientation. The 22 substituted positions are not preferentially in contact in this system: their mean minimum distance to lipid, 5.19 Å, is indistinguishable from the 5.21 Å of the remaining 135 positions. In the fully embedded state, which face of the bundle meets lipid is therefore set by the geometry of the insertion rather than by which positions were converted.

**Figure 5.**
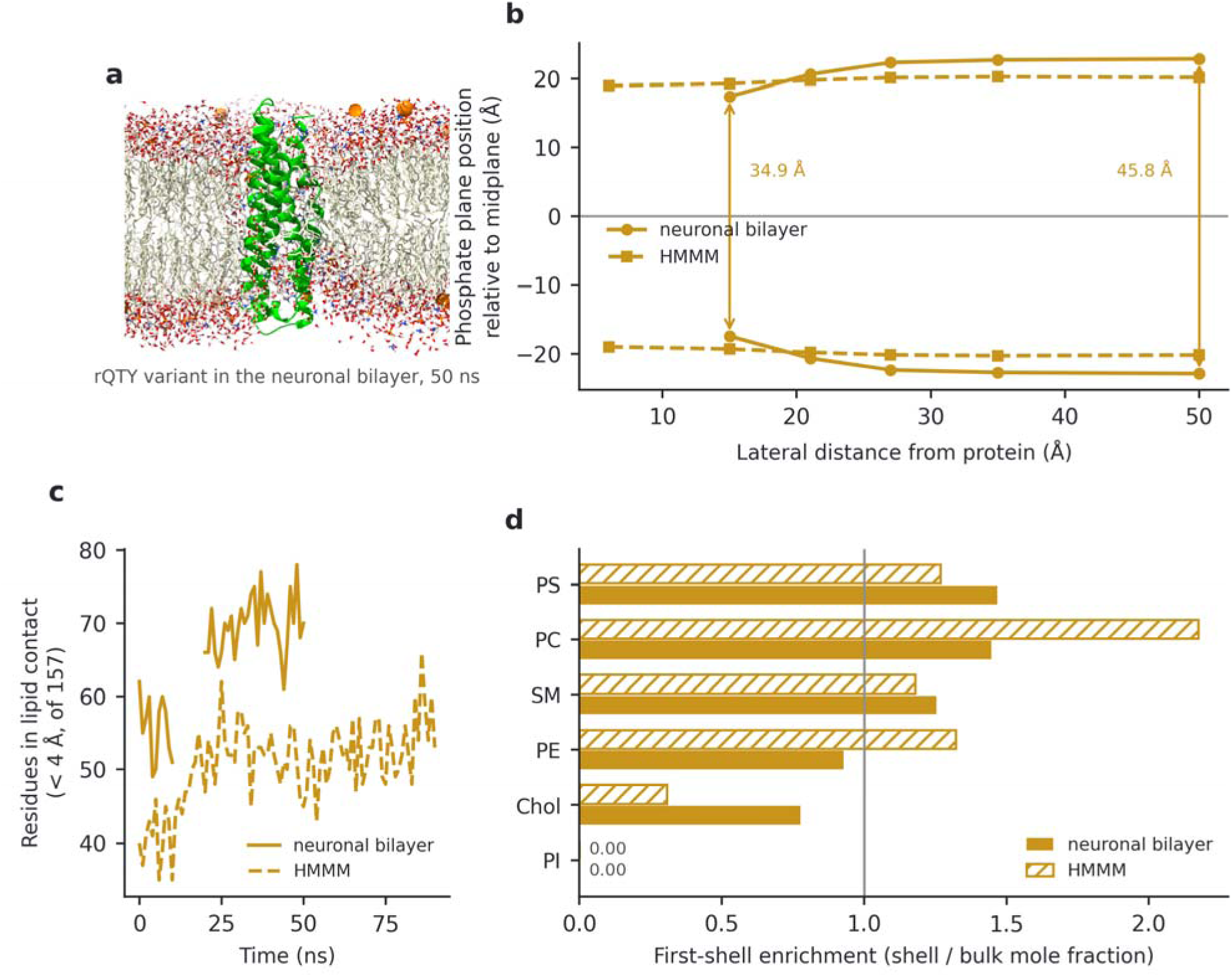
Membrane engagement of the embedded rQTY variant. (a) Snapshot of the variant in the six-component neuronal bilayer at 50 ns; the protein is drawn as a green ribbon, lipid acyl chains as pale cream sticks, headgroup and water oxygens in red, nitrogens in blue and headgroup phosphorus in small orange sticks; the large orange spheres in the aqueous phase are potassium ions. (b) Mean position of the upper and lower leaflet phosphate planes as a function of lateral distance from the protein centre of mass, averaged over all production frames, the one quantity in this study not restricted to the second half of each trajectory, because annuli close to the protein are sparsely occupied in any single frame. Distances are plotted at the centre of each annulus. Comparing the far field (40–60 Å) with the innermost annulus that both systems populate (12–18 Å), the phosphate-to-phosphate distance falls from 45.8 Å to 34.9 Å in the neuronal bilayer but only from 40.4 Å to 38.6 Å in the HMMM. The 0–12 Å annulus of the bilayer run contains too few phosphorus atoms to average and is omitted; the corresponding HMMM point (38.0 Å) is retained. (c) Number of residues within 4 Å of a lipid heavy atom as a function of time. (d) First-shell lipid enrichment, calculated as the mole fraction of species within 5 Å of the protein divided by its bulk mole fraction, over the second half of each each trajectory; the vertical line marks no enrichment. Cholesterol is depleted in both systems; phosphatidylinositol, present at six and four molecules respectively, is not observed in the shell of either. PS, phosphatidylserine; PE, phosphatidylethanolamine; PC, phosphatidylcholine; phosphatidylinositol; Chol, cholesterol; SM, sphingomyelin.

### Local Bilayer Deformation and Lipid Shell Composition

The most visible consequence of embedding is a graded deformation of the membrane around the protein. Measuring the phosphate-to-phosphate distance in annular bins centred on the protein gives 45.8 Å at 40–60 Å lateral distance, 45.5 Å at 30–40 Å, 44.7 Å at 24–30 Å, 41.3 Å at 18–24 Å and 34.9 Å at 12–18 Å. The bilayer is thus 10.9 Å thinner in the innermost populated annulus than in the far field of the same trajectory, with the perturbation decaying over roughly 20–30 Å (Fig. 5b). The annular profile is the one quantity here averaged over all production frames rather than the second half, because annuli close to the protein are sparsely occupied in any single frame. The reference is the far-field annulus of the same trajectory rather than a separately simulated protein-free bilayer, which was not run. Accompanying the deformation, 85.1 ± 4.6 water molecules occupy the hydrophobic core region (within 10 Å of the midplane and 25 Å laterally of the protein), indicating a hydrated defect tracking the protein through the bilayer rather than a cleanly sealed hydrophobic seam.

The lipid shell that forms around the protein is only mildly distinct from the bulk membrane. Counting lipids with any heavy atom within 5 Å of the protein over the equilibrated trajectory and comparing with bulk mole fractions gives enrichment factors of 1.47 for POPS, 1.45 for POPC, 1.25 for sphingomyelin and 0.93 for POPE, against 0.78 for cholesterol; phosphatidylinositol, of which the system contains six molecules, is not observed in the shell at any point (Fig. 5d). The clearest feature is the depletion of cholesterol, which makes up 45.5 mol% of the membrane but only 35% of the 39 lipids in the first shell. Phosphatidylserine is the most enriched species, but only marginally ahead of phosphatidylcholine, so the shell is not selectively anionic on this measure. Electrostatics give a clearer signal than composition does. The construct as a whole is net anionic (18 Arg and 7 Lys against 7 Asp and 20 Glu, net −2), so any preference for anionic lipid is not a consequence of overall charge; it is consistent instead with the basic surface patches of the N-terminal domain, of which the receptor-binding stretch 136–150 is the largest at net +7 [10, 11]. Basic side chains engage the headgroup region broadly: 23.5 ± 3.6 contacts within 4 Å are formed between arginine and lysine nitrogen atoms and the phosphate oxygens of all phospholipid species, occupying 15.8 ± 2.3 of the 61 basic nitrogen atoms at any instant. Counted over the same atoms of the anionic POPS and POPI species alone, 5.0 ± 2.1 of those contacts are with anionic lipid: 21% of basic-nitrogen contacts directed at species making up 15% of the phospholipid, a 1.4-fold anionic bias. That bias is the more robust of the two observations, because it is a per-frame count over 61 donor atoms rather than a mole fraction over a shell of 39 lipids. Lipid shells equilibrate more slowly than protein conformation, and with 24 sphingomyelin and six phosphatidylinositol molecules in a single 50 ns run the composition of the shell is the least converged quantity reported here; we therefore read it as a weak trend and do not interpret the ordering of the phospholipids further. The model bilayer is also symmetric and does not represent leaflet asymmetry, so these preferences describe the composition simulated here rather than a specific face of a neuronal membrane.

### Comparison with a Highly Mobile Membrane Mimetic

The same variant was simulated for 90 ns in a highly mobile membrane mimetic of matched headgroup composition (160 cholesterol, 80 POPE, 68 POPC, 24 POPS, 16 sphingomyelin, 4 POPI) in which the acyl chains are replaced by 704 1,1-dichloroethane molecules [36], giving a membrane with the same interfacial chemistry but far greater lateral fluidity. The behaviour of the protein is consistent with the full bilayer. It remains embedded, with the fraction of heavy atoms between the phosphate planes rising from 71% to 76%; the bundle Cα RMSD plateaus at 1.81 ± 0.14 Å; the radius of gyration is 19.33 ± 0.08 Å, within 0.1 Å of the full-bilayer value; and helix content is 87.7 ± 1.4%, with every bundle helix above 93% and H0 at 75.8% (Fig. 4, Table 3, Supplementary Table S3). That two membrane models differing substantially in acyl-region representation give the same fold, the same dimensions and the same helicity argues that these are properties of the construct rather than of the particular lipid parameterisation.

Three quantities differ, and all three differ in the direction expected from the way the mimetic is built. The bilayer deformation is much smaller in the mimetic, where the phosphate-to-phosphate distance falls only from 40.4 Å in the far field to 38.6 Å at 12–18 Å, a thinning of 1.8 Å on the unrounded values, against 10.9 Å in the full bilayer (Fig. 5b). A membrane whose interior is a small-molecule solvent accommodates an inserted protein by local rearrangement rather than by leaflet distortion. Protein–lipid contacts are correspondingly fewer, 294 ± 33, involving 52.5 ± 4.4 residues (Fig. 5c), since much of the buried protein surface faces dichloroethane rather than lipid. Apolar SASA is highest of the three systems at 4470 ± 88 Å², against 4047 ± 96 Å² in the full bilayer; the native value in water, 2226 ± 94 Å², is lower for the trivial reason that the native sequence carries 22 fewer apolar residues, and is listed in Table 3 for completeness rather than as a comparison.

Shell composition does not transfer between the two membrane models. In the mimetic the enrichment factors are 2.17 for POPC, 1.32 for POPE, 1.27 for POPS and 1.18 for sphingomyelin, against 0.31 for cholesterol, with phosphatidylinositol again absent (Fig. 5d). Cholesterol is depleted in both systems, and more strongly here, but the ordering of the phospholipids differs and phosphatidylcholine rather than phosphatidylserine dominates the mimetic shell. The mimetic is designed to give the acyl region greater lateral mobility, which should let the lipid shell equilibrate faster than in the full bilayer; we did not measure lateral diffusion or shell exchange, so that remains an expectation rather than a demonstration. Taken together, the one compositional effect the two models agree on is the exclusion of cholesterol from the first shell. Everything else differs between them, which is what a pair of unreplicated runs of 50 and 90 ns should be expected to deliver for a quantity that equilibrates on a longer timescale than either.

### Implications for ApoE Lipid Handling

These observations bear on why ApoE lipid handling is isoform- and lipidation-sensitive in the first place. The pathological phenotypes associated with ApoE4, including impaired amyloid-β clearance [7], exacerbated tau-mediated neurodegeneration [8] and lipid-droplet accumulation in microglia [9], all involve the protein operating at a lipid interface whose composition is itself disease-modified. A construct whose interfacial surface chemistry can be set by a defined substitution set, and which sits in a lipid shell from which cholesterol is excluded, offers a way to probe that dependence directly rather than through the abundance or receptor-binding routes that current ApoE-directed strategies use [4, 16, 18]. The design was carried out on the ApoE3 sequence, and because no substitution falls at residue 112 or 158 the same rule transfers unchanged to ApoE2 and ApoE4, so the isoform dependence of these phenotypes can be addressed with the identical substitution set on each background.

### Scope and Limitations of the Present Dataset

Three features of the dataset bound every statement above. Each membrane system was run once, so the standard deviations quoted are the spread of a single equilibrated time series (22 frames for the bilayer run, 50 for the mimetic) and not uncertainties of a mean over independent replicates; values are reported to the precision at which they were computed, but small differences between systems should not be read as resolved. No native control was simulated in either membrane, and no protein-free bilayer of the same composition was run, so the membrane observations characterise the variant against the far field of its own trajectory rather than against a native or an unperturbed reference. The bundle was placed inside the membrane at the build stage rather than reaching it from solution, which means these simulations test whether an embedded rQTY bundle is stable, not whether it partitions; nothing here measures affinity, and the comparison between native and variant remains one of sequence, hydropathy and predicted structure. Within those bounds the lipid-shell compositions are the least converged quantity reported, being drawn from 24 sphingomyelin and six phosphatidylinositol molecules in a single 50 ns run, and the two membrane models agree only on the depletion of cholesterol. The electrostatic contact counts, which are per-frame sums over 61 donor atoms rather than mole fractions over a few dozen lipids, are correspondingly better determined and carry the anionic-preference argument. A fourth bound applies to the design rather than to the trajectories: the substitution set is fixed by the secondary-structure assignment of one model, and while helix interiors are assigned identically by two experimental structures, three helix termini are not, so Q81, Q163 and Q21 are boundary calls rather than settled ones (Supplementary Tables S4 and S5).

A fifth bound is external to the dataset and concerns what a structure predictor can be asked to show. Systematic tests report that AlphaFold predictions are largely insensitive to substitution: where an evolutionary profile is available, nearly every sequence carrying 10% mutations passes any pLDDT threshold and more than 60% still pass pLDDT ≥ 70 at 30% mutations [40], while predicted coordinates can remain effectively invariant to the mutation of up to 40% of residues and AlphaFold3 confidence identifies the most accurate of several models only 16 to 35% of the time [41]. At 14.0% substitution the present design lies inside that insensitive regime, so the 0.81 Å mean agreement between the native and variant ensembles is a necessary condition for fold retention rather than evidence of it, and is reported here as such. Three elements of the analysis do not depend on the predictor discriminating between the two sequences. The native model from which the substitution set was read reproduces the wild-type crystal structure to 0.85 Å over 142 Cα atoms, so the secondary-structure assignment the rule was applied to is anchored to experiment and not to the prediction alone. The molecular dynamics uses a fixed empirical force field with no learned component, so the stability of the embedded variant over 50 and 90 ns is independent of the prediction entirely. The hydropathy, hydrophobic-moment and solvent-accessibility changes are computed from the sequence and from coordinates directly, and are large relative to any plausible prediction error. The AlphaFold3 ensembles are therefore used to establish that the conversion introduces no predicted fold change, which is what the comparison can support, while the affirmative statements rest on the crystallographic control and on the simulations.

## Conclusion

We set out to test whether the reverse-QTY code can be used as a controlled, residue-resolved dial on the surface chemistry of a water-soluble helical bundle, and whether the resulting construct is compatible with a lipid environment, using the N-terminal domain of human apolipoprotein E as the test case. The design rule was minimal: within mature residues 11–167, every glutamine, threonine and tyrosine lying inside an α-helix was converted to leucine, valine or phenylalanine, and every such residue in a turn or loop was left alone. That single criterion produced 22 substitutions across 14.0% of the segment and left 86.0% of the sequence native. It also produced, without any further intervention, two boundary conditions that a rational design would have had to impose by hand: no substitution falls between residues 124 and 155, so the LDLR recognition region is untouched, and the conversion stops at the hinge, so the C-terminal high-affinity lipid-binding domain is unaltered.

The structural consequence of that conversion is contained. Across five AlphaFold3 models per sequence, the four-helix bundle of the variant superimposes on the native bundle at 0.81 Å mean Cα RMSD, no cross-ensemble pair exceeding 1.02 Å, with indistinguishable helix boundaries and helix content. The heterogeneity of the variant ensemble resolves to the N-terminal arm (residues 11–24), a low-confidence, mobile element in the native sequence as well. The chemistry of the accessible surface meanwhile changes substantially, apolar solvent-accessible area rising by 59% at unchanged total area and unchanged burial of the substituted positions, and it does so while raising rather than flattening the hydrophobic moment of five of the six helical segments. The variant remains amphipathic, which is the property that governs how apolipoprotein helices behave at an interface; the single segment that loses facial asymmetry is the one the conversion makes most hydrophobic.

In molecular dynamics, the native segment holds its fold in water over 100 ns, so the isolated fragment does not need the C-terminal domain to stay folded on that timescale. The rQTY variant, built into a six-component neuronal bilayer and into a matched highly mobile membrane mimetic, remains folded and essentially fully helical in both (helix content 86.4 ± 1.2% and 87.7 ± 1.4%; bundle RMSD 1.94 ± 0.12 Å and 1.81 ± 0.14 Å). That two membrane representations differing substantially in how the acyl region is modelled agree on fold, dimensions and helicity argues that the stability is a property of the construct rather than of one lipid model. Around the embedded variant the bilayer is 10.9 Å thinner in the innermost populated annulus than in the far field. The first lipid shell is mildly biased rather than strongly selective: cholesterol is depleted in both membranes, and this is the only compositional effect the two models agree on, while a modest preference for anionic lipid shows up more clearly in the basic-nitrogen contact counts than in the shell mole fractions. With a single unreplicated run of each system, shell composition remains the least converged quantity in the study.

These results indicate that a geometry-preserving chemical conversion of helical Q, T and Y positions leaves a soluble apolipoprotein bundle able to occupy a lipid bilayer, once embedded, with its fold, its helicity and its receptor-binding sequence intact. Two further steps would extend that conclusion. Simulating the native segment in the identical bilayer and mimetic, in replicate and at matched length, would convert the present sequence- and structure-level comparison into a direct comparison of membrane behaviour; and a free-energy calculation of partitioning from bulk solvent would address affinity itself, which the embedded starting configuration used here is not designed to measure. Experimental validation follows the same logic: circular dichroism to confirm helicity, and tryptophan fluorescence or liposome flotation to quantify binding of native and variant under identical lipid conditions. The segment carries four tryptophans, at positions 20, 26, 34 and 39, none of which the rQTY rule alters, so the fluorescence readout would be identical in composition between the two constructs and would report on the arm and on H1.

Beyond ApoE, the result speaks to the symmetry of the QTY code itself. The forward direction has been used repeatedly to make hydrophobic membrane proteins water-soluble while preserving their fold; here the reverse direction, applied for the first time to a soluble helical bundle rather than to build an amphiphile *de novo*, preserves the predicted and simulated fold just as faithfully while moving the surface chemistry the other way. The same substitution pairs therefore appear to operate as a bidirectional, residue-resolved control on the solvent preference of an α-helical protein, with the secondary structure supplying the selection rule and the fold absorbing the change. For proteins whose biology is executed at a lipid interface, above all the exchangeable apolipoproteins but also the wider class of amphipathic-helix membrane sensors and remodellers, this offers a way to treat interfacial affinity as a designable parameter rather than a fixed property, and so to ask which of their functions actually depend on it. Combining such designed variants with dynamics-aware evolutionary analysis may further indicate which positions tolerate a change of side-chain chemistry and which are constrained by motion rather than by packing [42].

## Materials and Methods

### Sequence Design

All residue numbering refers to mature human ApoE (UniProt P02649, residues 19–317 of the precursor), which is the ApoE3 isoform (Cys112, Arg158). Both polymorphic positions lie inside the designed segment, and neither is a glutamine, threonine or tyrosine, so the rQTY rule leaves them unchanged and transfers without modification to the ApoE2 and ApoE4 backgrounds. The design target was the segment spanning residues 11–167, which contains the complete four-helix bundle of the N-terminal domain and stops short of the interdomain hinge. Secondary structure was assigned with the Dictionary of Secondary Structure of Proteins (DSSP) algorithm [43] on the top-ranked AlphaFold3 [24] model of the complete 299-residue mature ApoE sequence, predicted on the AlphaFold Server with model seed 160824601 (pTM 0.53, ranking score 0.75 for the full-length chain). Confidence in that model is strongly region-dependent, as expected for a two-domain protein joined by a flexible hinge: mean per-residue pLDDT is 72.5 over the whole chain, but 85.7 over the segment used for the design (residues 11–167) and 89.2 over the four-helix bundle itself (residues 25–167), against 61.5 over the C-terminal domain (residues 192–299). The region from which the design was taken is therefore well determined in the model that defined it. The reverse-QTY code was then applied under a single rule: every glutamine, threonine and tyrosine residue whose DSSP assignment was helical (H, G or I) was converted according to the rQTY pairings Q→L, T→V and Y→F, and every glutamine, threonine or tyrosine outside a helix was left unchanged. Because the forward QTY code maps both isoleucine and valine onto threonine, the reverse assignment of threonine is not unique; T→V was chosen over T→I because the T V exchange is the one we have found to be evolutionarily coupled in α-helical membrane proteins [26, 33], and the same choice is applied to all four threonine positions. No criterion of burial, conservation or helical face was used. The helices assigned in that model span residues 11–19, 25–41, 45–52, 55–81, 87–124, 131–198 and 202–288, of which the first six lie wholly or partly within the design segment. Throughout, these six are labelled H0 (11–19), H1 (25–41), H1b (45–52), H2 (55– 81), H3 (87–124) and H4 (131–167), H4 being the 131–198 helix truncated at the design boundary; H1– H4 are the helices of the bundle proper, H0 is the helical part of the N-terminal arm and H1b a short connecting helix. This produced 22 substitutions (14 Q→L, 4 T→V, 4 Y→F), listed in full in Table 1.

Because the substitution set follows from a secondary-structure assignment, it inherits whatever uncertainty that assignment carries. The rule was therefore reapplied using experimental structures of the same protein in place of the predicted model: the crystal structure of the wild-type ApoE3 N-terminal domain (PDB 1LPE, 2.25 Å, mature residues 23–166) [10] and the solution NMR structure of full-length ApoE3 (PDB 2L7B, 20 conformers) [13], whose designed segment was confirmed to carry the native sequence, the monomerising substitutions of that construct lying entirely outside residues 11–167. DSSP was computed with the same implementation in all three cases, and for the NMR ensemble a position was taken as helical when at least half of the 20 conformers assigned H, G or I. Supplementary Tables S4 and S5 give the resulting helix boundaries and the position-by-position comparison.

### Structure Prediction

Native and rQTY sequences were submitted separately to the AlphaFold3 server [24] using the alphafoldserver dialect (version 1) with structure templates enabled for both jobs. Single model seeds were used for each job (481923589 for the native segment, 693416453 for the variant), and five models were generated per sequence. Template search was run by the server for both sequences; the identity of the templates retrieved is not recorded in the downloaded output, and because the variant differs from the native sequence at 22 positions the two jobs need not have drawn on the same templates. This is a possible contributor to the difference in global confidence between the two ensembles, and is one reason the comparison in Results is drawn on region-resolved rather than global metrics. Per-model confidence metrics (predicted TM-score, pTM; ranking score; fraction disordered; clash flag), per-atom predicted local distance difference test (pLDDT) values and the predicted aligned error matrix were taken directly from the server output. For the two segment jobs the server returned a ranking score equal to pTM, 0.85 for the native segment and 0.79 for the variant; for the full-length job the ranking score (0.75) exceeds pTM (0.53) because of the disorder term the server includes in its ranking. The top-ranked model of each sequence was used as the starting structure for molecular dynamics.

### Hydropathy and Amphipathicity Analysis

Grand average of hydropathicity (GRAVY) was computed on the Kyte–Doolittle scale [35] over the whole segment and in a nine-residue sliding window. Mean hydrophobicity H and mean hydrophobic moment μH were computed on the Eisenberg consensus scale [37], with the moment evaluated at 100° per residue for an ideal α-helix [37]; both were resolved by helical segment using the DSSP boundaries above.

### System Construction

All simulation systems were built with CHARMM-GUI [44], using Solution Builder for the aqueous system, Membrane Builder [45] for the explicit bilayer and HMMM Builder [46] for the membrane mimetic. The CHARMM36m force field was used for protein [47], CHARMM36 for lipids [48], and the CHARMM-modified TIP3P model for water [49, 50]. All systems were neutralised and brought to 0.15 M KCl.

The aqueous system contained the native segment, 21,113 water molecules, 62 K and 60 Cl in a cubic box of 8.68 nm per side (66,051 atoms). The explicit bilayer contained the rQTY segment in a six-component membrane chosen to approximate the lipid classes of a neuronal plasma membrane (240 cholesterol, 120 POPE, 102 POPC, 36 POPS, 24 sphingomyelin and 6 POPI, 528 lipids in total; 45.5 mol% cholesterol). The two leaflets were built with the same composition, so the model does not represent the leaflet asymmetry of a plasma membrane. The system also contained 31,947 water molecules, 131 K and 87 Cl in a box of 11.03 × 11.03 × 12.12 nm (153,551 atoms). The membrane mimetic contained the rQTY segment in a bilayer of matched headgroup composition (160 cholesterol, 80 POPE, 68 POPC, 24 POPS, 16 sphingomyelin, 4 POPI) whose acyl chains were truncated at the sixth carbon and replaced by 704 1,1-dichloroethane molecules parameterised with CGenFF as distributed with the CHARMM-GUI HMMM Builder, with 23,754 water molecules, 95 K and 65 Cl in a box of 9.40 × 9.40 × 11.91 nm (103,936 atoms). Complete compositions of all three systems are given in Supplementary Table S2.

The designed segment is an internal fragment of the mature chain and was built with free charged termini (protonated N-terminus, deprotonated C-terminus) rather than with neutral caps. In both membrane systems the protein was inserted into the bilayer at the build stage rather than placed in bulk solvent, so that the simulations test the stability of the membrane-embedded state rather than spontaneous partitioning from solution. At the start of production, 75% (explicit bilayer) and 71% (mimetic) of protein heavy atoms lay between the two leaflet phosphate planes.

### Simulation Protocol

All simulations were run in GROMACS 2024.4 [51] in mixed precision, in a cloud-based execution environment whose performance for systems of this size we have benchmarked previously [52]. Each system was energy-minimised by steepest descent to a maximum force below 1000 kJ mol ¹ nm ¹ (at most 5000 steps). Membrane systems were then equilibrated through the standard six-stage CHARMM-GUI protocol totalling 1.875 ns (three 125 ps stages at a 1 fs timestep followed by three 500 ps stages at 2 fs), with positional restraints on protein backbone and side-chain atoms released progressively from 4000 and 2000 to 50 and 0 kJ mol ¹ nm ², lipid restraints from 1000 to 0 kJ mol ¹ nm ², and lipid dihedral restraints from 1000 to 0 kJ mol ¹ rad ². The aqueous system was equilibrated for 125 ps at a 1 fs timestep with backbone and side-chain restraints of 400 and 40 kJ mol ¹ nm ².

Production runs used a 2 fs timestep with bonds to hydrogen constrained by LINCS [53], a Verlet neighbour list updated every 20 steps, van der Waals interactions cut off at 1.2 nm with a force switch from 1.0 nm, and electrostatics by particle-mesh Ewald [54] with a 1.2 nm real-space cutoff. Temperature was held at 303.15 K with the velocity-rescaling thermostat [55] (τ = 1 ps) applied to solute, membrane and solvent groups separately (solute and solvent only in the aqueous system), and pressure at 1 bar with the stochastic cell-rescaling barostat [56] (τ = 5 ps, compressibility 4.5 × 10 bar ¹), semi-isotropically for the membrane systems and isotropically for the aqueous system. In the mimetic, positional restraints of 200 kJ mol ¹ nm ² were retained during production on the terminal carbon atoms of the truncated acyl chains along the bilayer normal only, as required by the HMMM model to hold the two leaflets apart while leaving lateral mobility unrestricted [36, 46]; no restraints were applied to the protein or to lipid headgroups. Coordinates were written every 1 ns, so the equilibrated averages reported here are taken over 55, 22 and 50 frames for the aqueous, bilayer and mimetic systems respectively. Each system was run once, and the standard deviations quoted throughout are the spread of that single equilibrated time series rather than standard errors over independent replicates. Production lengths were 100 ns for the native segment in water, 50 ns for the rQTY variant in the explicit bilayer and 90 ns for the rQTY variant in the mimetic. Each was run as consecutive 10 ns segments of one continuous trajectory, not as independent replicates; the frame counts in Supplementary Table S2 include the duplicated frame at each segment boundary. In the bilayer run the coordinate file for the 10–20 ns segment was overwritten by an interrupted re-run and is unavailable, so analysis of that system uses the 44 retained frames, which span 0–10 and 20–50 ns; the equilibrated averages fall entirely after 30 ns and are unaffected (Supplementary Note S2).

### Trajectory Analysis

Trajectories were analysed with MDAnalysis 2.10.0 [57, 58] and MDTraj 1.11.1 [59], within the analysis framework we developed for larger all-atom studies of neuronal transporter assembly in explicit membranes [60]. Before any structural analysis the protein was made whole across periodic boundaries by a sequential minimum-image walk along the atom ordering (Supplementary Note S2). Cα RMSD was computed after Kabsch superposition, either on all 157 residues or on the bundle alone (residues 25– 167), as stated. Root-mean-square fluctuation (RMSF) was computed over the second half of each trajectory after superposition on the bundle. Inter-helical separations are distances between the Cα centroids of the helices concerned, averaged over the five models of an ensemble. Secondary structure was assigned frame by frame with DSSP [43] as implemented in MDTraj, helix content being the fraction of residues assigned H, G or I. Solvent-accessible surface area (SASA) was computed by the Shrake–Rupley algorithm [38] with a 1.4 Å probe radius and 960 sphere points per atom; apolar SASA is the sum over Ala, Val, Leu, Ile, Phe, Met, Trp, Pro and Cys. Relative solvent accessibility (RSA) used the theoretical maximum accessibilities of Tien et al. [39].

For the membrane systems, the bilayer midplane was defined by the mean z coordinate of all lipid phosphorus atoms in each frame, and the two leaflet phosphate planes by the mean z of the phosphorus atoms above and below it. Local membrane deformation was quantified by binning phosphorus atoms into annuli of 0–12, 12–18, 18–24, 24–30, 30–40 and 40–60 Å lateral distance from the protein centre of mass and computing the leaflet separation within each annulus; annuli containing fewer than three phosphorus atoms in a given leaflet were excluded from that frame. Because annuli close to the protein are sparsely occupied in any single frame, the annular profile alone is averaged over all production frames rather than over the second half. Protein–lipid contacts were counted as heavy-atom pairs within 4 Å, and a residue was counted as in contact if any of its heavy atoms was within 4 Å of a lipid heavy atom. First-shell lipid composition was defined by lipid molecules with any heavy atom within 5 Å of the protein, and enrichment as the ratio of a species’ mole fraction in that shell to its bulk mole fraction. Core hydration was counted as water oxygens within 10 Å of the midplane and 25 Å laterally of the protein centre of mass. Contacts between basic side chains and lipid headgroups were counted in two ways, in both cases as atom pairs within 4 Å of arginine NE/NH1/NH2 or lysine NZ nitrogens: first against the phosphate oxygens (O11, O12, O13, O14) of all phospholipid species, and second against the same four atoms of the anionic POPS and POPI species alone, so that the second count is a strict subset of the first. Unless stated otherwise, all reported means and standard deviations are over the second half of each production run.

## Statements and Declarations

### Supplementary Information

Supplementary Notes S1 and S2, Supplementary Tables S1–S5 and Supplementary Figs. S1 and S2 are provided as a separate file (Supplementary_Information.docx).

### Author Contributions

T.K. and A.K. contributed equally to this work. Both authors designed the study, carried out the computational work and the analysis, wrote the manuscript, and read and approved the final version.

### Data Availability

The native and reverse-QTY sequences of the designed segment are given in full in Supplementary Note S1. The experimental structures used for comparison are available from the RCSB Protein Data Bank (https://www.rcsb.org) under accessions 1LPE and 2L7B. The AlphaFold3 model of full-length mature ApoE from which the secondary-structure assignment was taken, the AlphaFold3 models of the native and reverse-QTY segments with their confidence files, the CHARMM-GUI starting coordinates and topologies and every GROMACS parameter file for all three simulated systems, the analysis outputs from which every reported value is drawn, and the Python scripts that perform the sequence, structure and trajectory analysis and generate every figure can be accessed at https://github.com/karagol-taner/apoe-reverse-qty. The deposited inputs are sufficient to repeat each simulation from the starting coordinates; the production trajectories themselves are too large for that repository and are available from the corresponding authors on request. There are no restrictions on availability.

### Ethics Approval

Ethics approval was not required for this computational study, which did not involve animal subjects, human participants or identifiable data.

### Consent to Participate

Not applicable. This computational study did not involve human participants.

### Consent to Publish

Not applicable. This computational study did not involve human participants.

### Declaration of Generative AI and AI-Assisted Technologies in the Manuscript Preparation Process

During the preparation of this work, the authors used Claude Opus 5 (Anthropic) for drafting and for streamlining analysis code. The authors reviewed and edited the output as needed and take full responsibility for the content of the published article.

