## Supplementary_Information for "The Apolipoprotein E N-Terminal Bundle Rendered Membrane-Compatible by Reverse-QTY Conversion Without Loss of Fold"

Running Title: Reverse-QTY ApoE N-Terminal Bundle

#### Supplementary Note S1. Sequences of the designed segment

Both sequences span residues 11–167 of mature human ApoE (UniProt P02649), 157 residues. Numbering at the left of each block is the position of the first residue on that line in mature ApoE numbering. The two sequences differ at 22 positions (14.0%), all listed in Supplementary Table S1.

**ApoE-N (native), residues 11–167**

```
 11  EPELRQQTEWQSGQRWELALGRFWDYLRWVQTLSEQVQEELLSSQVTQELRALMDETMKE
 71  LKAYKSELEEQLTPVAEETRARLSKELQAAQARLGADMEDVCGRLVQYRGEVQAMLGQST
131  EELRVRLASHLRKLRKRLLRDADDLQKRLAVYQAGAR
```

**ApoE-N rQTY variant, residues 11–167**

```
 11  EPELRLLVEWQSGQRWELALGRFWDFLRWVLTLSELVLEELLSSLVVLELRALMDEVMKE
 71  LKAFKSELEELLTPVAEEVRARLSKELLAALARLGADMEDVCGRLVLFRGEVLAMLGQST
131  EELRVRLASHLRKLRKRLLRDADDLLKRLAVFLAGAR
```

### Supplementary Table S1. Every glutamine, threonine and tyrosine residue in the designed segment

The design rule converts a residue if and only if its DSSP assignment is helical. All 22 converted positions carry DSSP code H; all six unconverted positions lie outside helices. DSSP codes are H, α-helix; T, hydrogen-bonded turn; S, bend; C, coil. RSA, relative solvent accessibility averaged over the five AlphaFold3 models of each sequence. En dashes mark entries that do not apply because the residue was not converted.

| Position | Residue | DSSP | Helix | Converted | rQTY residue | RSA native (%) | RSA rQTY (%) |
|---|---|---|---|---|---|---|---|
| 16 | Q16 | H | H0 | yes | L16 | 22.3 | 34.3 |
| 17 | Q17 | H | H0 | yes | L17 | 18.4 | 32.7 |
| 18 | T18 | H | H0 | yes | V18 | 49.8 | 48.8 |
| 21 | Q21 | C | – | no | – | 59.5 | 50.1 |
| 24 | Q24 | C | – | no | – | 31.7 | 26.7 |
| 36 | Y36 | H | H1 | yes | F36 | 14.6 | 16.7 |
| 41 | Q41 | H | H1 | yes | L41 | 11.3 | 19.8 |
| 42 | T42 | T | – | no | – | 51.4 | 49.3 |
| 46 | Q46 | H | H1b | yes | L46 | 46.0 | 45.0 |
| 48 | Q48 | H | H1b | yes | L48 | 20.6 | 18.2 |
| 55 | Q55 | H | H2 | yes | L55 | 42.9 | 47.5 |
| 57 | T57 | H | H2 | yes | V57 | 12.8 | 10.7 |
| 58 | Q58 | H | H2 | yes | L58 | 51.5 | 56.1 |
| 67 | T67 | H | H2 | yes | V67 | 0.0 | 0.1 |
| 74 | Y74 | H | H2 | yes | F74 | 10.3 | 7.9 |
| 81 | Q81 | H | H2 | yes | L81 | 55.8 | 33.5 |
| 83 | T83 | C | – | no | – | 28.5 | 55.8 |
| 89 | T89 | H | H3 | yes | V89 | 31.0 | 34.0 |
| 98 | Q98 | H | H3 | yes | L98 | 50.4 | 47.6 |
| 101 | Q101 | H | H3 | yes | L101 | 10.8 | 5.4 |
| 117 | Q117 | H | H3 | yes | L117 | 42.3 | 48.7 |
| 118 | Y118 | H | H3 | yes | F118 | 1.9 | 2.2 |
| 123 | Q123 | H | H3 | yes | L123 | 55.4 | 51.9 |
| 128 | Q128 | S | – | no | – | 54.2 | 54.9 |
| 130 | T130 | C | – | no | – | 14.2 | 25.9 |
| 156 | Q156 | H | H4 | yes | L156 | 18.2 | 25.9 |
| 162 | Y162 | H | H4 | yes | F162 | 5.0 | 1.1 |
| 163 | Q163 | H | H4 | yes | L163 | 26.3 | 25.6 |

### Supplementary Table S2. Composition of the three simulation systems

En dashes mark components absent from a given system. All systems were neutralised and brought to 0.15 M KCl. DCLE, 1,1-dichloroethane, which replaces the truncated acyl chains in the highly mobile membrane mimetic. Each production run was carried out as consecutive 10 ns segments of one continuous trajectory; frame counts include the duplicated frame at each segment boundary, which is why they slightly exceed the nominal length in nanoseconds. The second figure in each pair is the number of frames entering the equilibrated averages.

| Component | Native, water | rQTY, neuronal bilayer | rQTY, HMMM |
|---|---|---|---|
| Protein (residues) | 157 | 157 | 157 |
| Cholesterol | – | 240 | 160 |
| POPE | – | 120 | 80 |
| POPC | – | 102 | 68 |
| POPS | – | 36 | 24 |
| Sphingomyelin (PSM) | – | 24 | 16 |
| POPI | – | 6 | 4 |
| 1,1-dichloroethane (DCLE) | – | – | 704 |
| Water (TIP3P) | 21,113 | 31,947 | 23,754 |
| $K^+$ / $Cl^-$ | 62 / 60 | 131 / 87 | 95 / 65 |
| Total atoms | 66,051 | 153,551 | 103,936 |

| | | | |
|---|---|---|---|
| Box (nm) | 8.68 × 8.68 × 8.68 | 11.03 × 11.03 × 12.12 | 9.40 × 9.40 × 11.91 |
| Barostat coupling | isotropic | semi-isotropic | semi-isotropic |
| Production (ns) | 100 (10 × 10) | 50 (44 frames retained) | 90 (9 × 10) |
| Frames written / analysed | 110 / 55 | 44 / 22 | 99 / 50 |

### Supplementary Table S3. Helicity of the bundle helices by segment (%)

Segments are the six DSSP helices of the designed region; H1–H4 are the helices of the bundle proper, H0 the helical part of the N-terminal arm and H1b a short connecting helix. AlphaFold3 columns indicate whether the segment is predicted helical in the top-ranked model of each sequence. Molecular dynamics columns give the percentage of frames in the second half of each trajectory in which residues of that segment are assigned H, G or I by DSSP. The connecting-loop and whole-segment rows are given so that the per-segment values reconcile with the whole-segment helix content quoted in Table 3: helix content is computed over all 157 residues, including those outside the reference helix boundaries.

| Segment | Residues | n | AlphaFold3 native | AlphaFold3 rQTY | MD native, water | MD rQTY, bilayer | MD rQTY, HMMM |
|---|---|---|---|---|---|---|---|
| H0 | 11–19 | 9 | helical | helical | 77.8 | 77.8 | 75.8 |
| H1 | 25–41 | 17 | helical | helical | 99.9 | 98.8 | 99.4 |
| H1b | 45–52 | 8 | helical | helical | 42.0 | 100.0 | 93.5 |
| H2 | 55–81 | 27 | helical | helical | 97.6 | 98.5 | 97.6 |
| H3 | 87–124 | 38 | helical | helical | 95.7 | 96.8 | 96.0 |
| H4 | 131–167 | 37 | helical | helical | 86.7 | 91.2 | 96.9 |
| Connecting loops | – | 21 | – | – | 18.4 | 31.6 | 37.0 |
| Whole segment | 11–167 | 157 | – | – | 80.3 | 86.4 | 87.7 |

### Supplementary Table S4. Helix boundaries from three structures

Helix boundaries of the designed segment from three structures. The design rule uses the DSSP assignment of the top-ranked AlphaFold3 model of full-length mature ApoE. The two experimental structures are the crystal structure of the wild-type ApoE3 N-terminal domain (PDB 1LPE, 2.25 Å) and the solution NMR structure of full-length ApoE3 (PDB 2L7B, 20 conformers), whose designed segment carries the native sequence. Boundaries are the limits of contiguous runs of at least four residues assigned H, G or I, computed with the same DSSP implementation in all three cases; for the NMR ensemble a residue counts as helical when at least half of the 20 conformers assign it so. 1LPE resolves mature residues 23 to 166, so H0 is absent from it and the C-terminal limit of H4 cannot be observed beyond residue 166; residues 163 to 166 are present in that model and are not helical. En dashes mark segments a structure does not resolve.

| Segment | AlphaFold3 (design model) | 1LPE, X-ray | 2L7B, NMR |
|---|---|---|---|
| H0 | 11–19 | – | 13–22 |
| H1 | 25–41 | 25–41 | 25–39 |
| H1b | 45–52 | 45–52 | 46–50 |
| H2 | 55–81 | 55–78 | 55–78 |
| H3 | 87–124 | 87–123 | 90–125 |
| H4 | 131–167 | 131–162 | 131–162 |

### Supplementary Table S5. Assignment at every Q, T and Y position from three structures

Secondary-structure assignment at every glutamine, threonine and tyrosine of the designed segment, from the same three structures. Converted records the decision taken in this work, which follows the AlphaFold3 column. For 2L7B the majority code is given with the number of the 20 conformers assigning it. Same decision records whether the rule, applied to the experimental assignment instead, would have made the same choice, and names the structures that dissent. 23 of the 28 positions are unanimous across every structure that resolves them. The five that are not lie at three helix termini: Q81 and Q163 were converted on a helical assignment that neither experimental structure supports, Q21 was retained although it is helical in all 20 NMR conformers, and Q41 and T89 are helical in the crystal structure and in the design model but not in the majority of NMR conformers. En dashes mark positions 1LPE does not resolve.

| Position | Residue | Helix | Converted | AlphaFold3 | 1LPE | 2L7B | Same decision |
|---|---|---|---|---|---|---|---|
| 16 | Q | H0 | yes | H | – | H (20/20) | yes |
| 17 | Q | H0 | yes | H | – | H (20/20) | yes |
| 18 | T | H0 | yes | H | – | H (20/20) | yes |
| 21 | Q | – | no | C | – | H (20/20) | no (2L7B) |
| 24 | Q | – | no | C | C | S (17/20) | yes |
| 36 | Y | H1 | yes | H | H | H (20/20) | yes |
| 41 | Q | H1 | yes | H | H | C (10/20) | no (2L7B) |
| 42 | T | – | no | T | T | C (19/20) | yes |
| 46 | Q | H1b | yes | H | H | H (20/20) | yes |
| 48 | Q | H1b | yes | H | H | H (20/20) | yes |
| 55 | Q | H2 | yes | H | H | H (20/20) | yes |
| 57 | T | H2 | yes | H | H | H (20/20) | yes |
| 58 | Q | H2 | yes | H | H | H (20/20) | yes |
| 67 | T | H2 | yes | H | H | H (20/20) | yes |
| 74 | Y | H2 | yes | H | H | H (20/20) | yes |
| 81 | Q | H2 | yes | H | S | S (13/20) | no (1LPE, 2L7B) |
| 83 | T | – | no | C | C | C (15/20) | yes |
| 89 | T | H3 | yes | H | H | T (13/20) | no (2L7B) |
| 98 | Q | H3 | yes | H | H | H (20/20) | yes |
| 101 | Q | H3 | yes | H | H | H (20/20) | yes |
| 117 | Q | H3 | yes | H | H | H (20/20) | yes |
| 118 | Y | H3 | yes | H | H | H (20/20) | yes |
| 123 | Q | H3 | yes | H | H | H (16/20) | yes |
| 128 | Q | – | no | S | S | C (18/20) | yes |
| 130 | T | – | no | C | C | S (20/20) | yes |
| 156 | Q | H4 | yes | H | H | H (20/20) | yes |
| 162 | Y | H4 | yes | H | H | H (14/20) | yes |
| 163 | Q | H4 | yes | H | T | T (18/20) | no (1LPE, 2L7B) |

### Supplementary Fig. S1

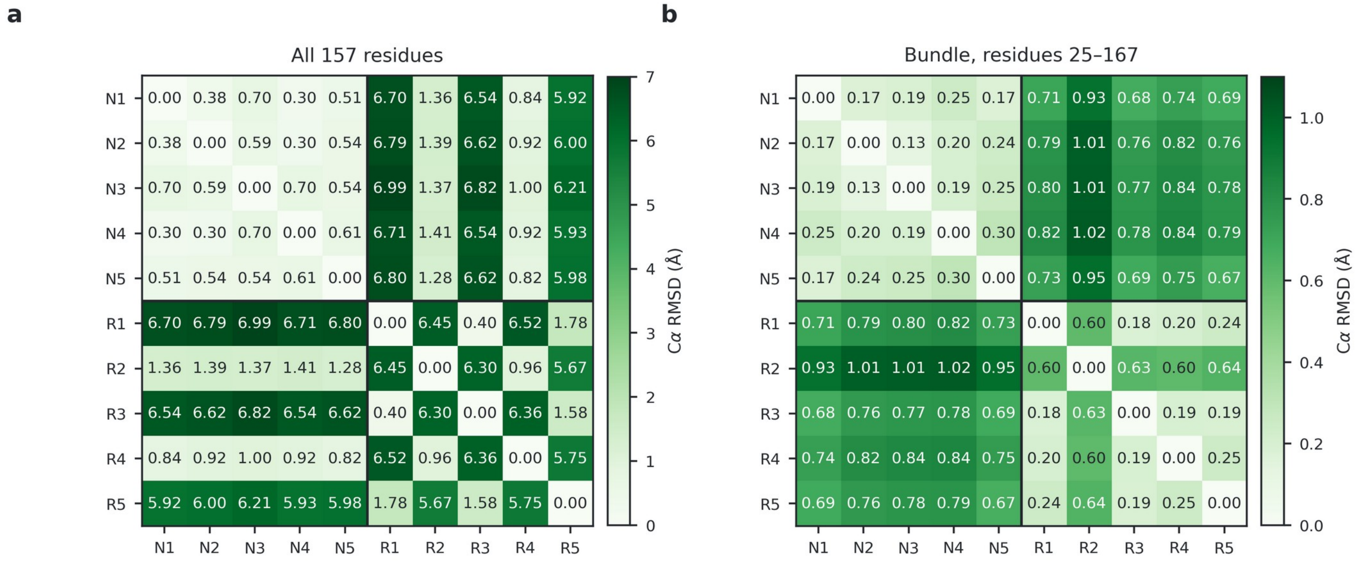


N1–N5, the five AlphaFold3 models of the native segment; R1–R5, the five models of the rQTY variant.

Fig. S1 Complete pairwise Cα RMSD matrices for the ten AlphaFold3 models. (a) Superposition over all 157 residues. The native models (N1–N5) form one tight cluster, while the rQTY models split into a native-like group (R2, R4) and a second group (R1, R3, R5) roughly 6 Å away. (b) The same ten models superposed over the bundle alone (residues 25–167). The block structure disappears: every native–variant pair lies between 0.67 and 1.02 Å, and the largest value anywhere in the matrix is 1.02 Å. Note the different colour scales.

**Supplementary Fig. S2**

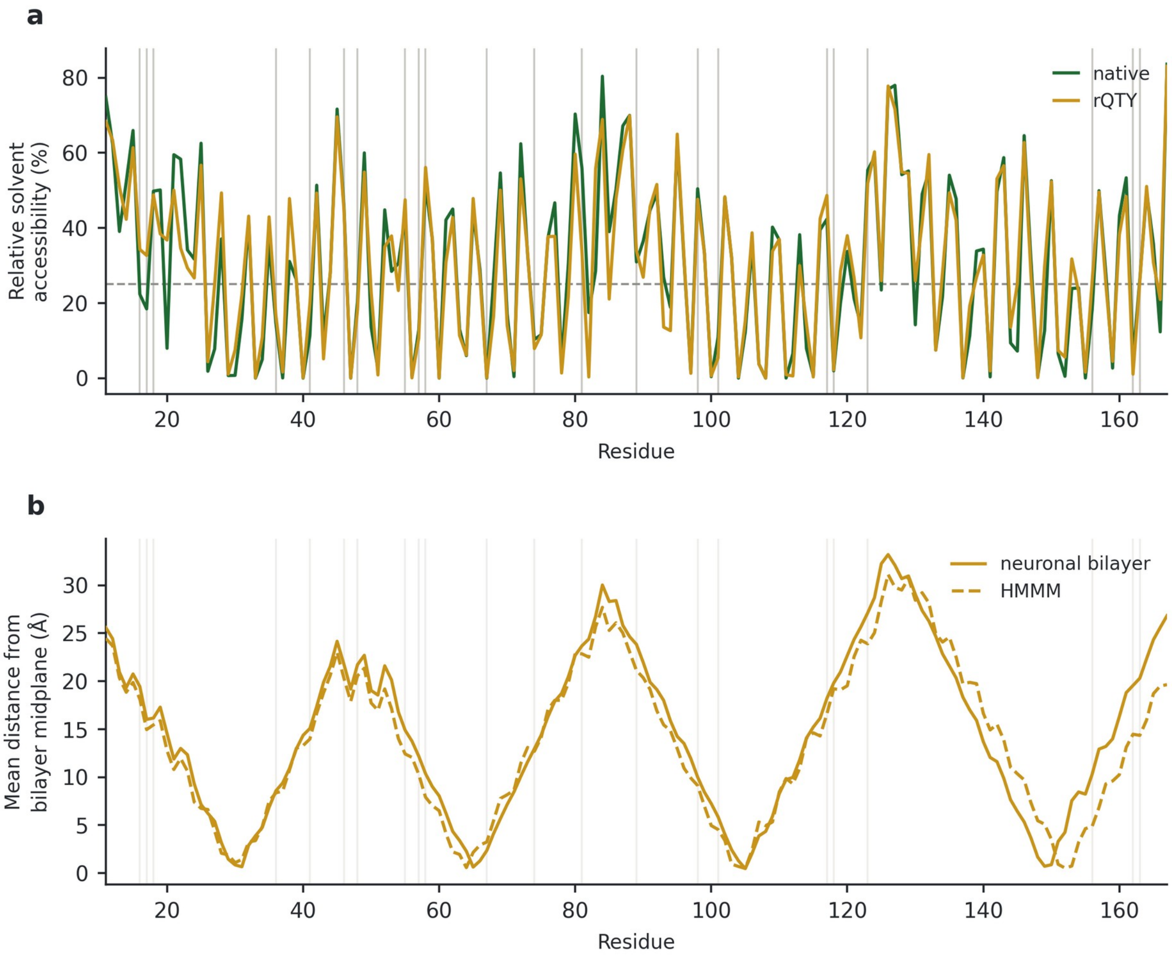


Fig. S2 Per-residue surface exposure and membrane depth. (a) Relative solvent accessibility of the native (green) and rQTY (gold) segments, averaged over the five AlphaFold3 models of each. Vertical grey lines mark the 22 substituted positions; the dashed horizontal line marks the 25% threshold used to classify a position as solvent-exposed. Exposure at the substituted positions is essentially unchanged by the substitution (mean 27.2% native, 27.9% rQTY). (b) Mean absolute distance of each residue from the bilayer midplane over the second half of each membrane trajectory, for the rQTY variant in the neuronal bilayer (solid) and in the HMMM (dashed); the faint vertical lines again mark the 22 substituted positions.

### Supplementary Note S2. Reproducibility

Every quantity reported in the main text was computed from the AlphaFold3 server output and the GROMACS production trajectories using MDAnalysis 2.10.0 and MDTraj 1.11.1. Two analysis details materially affect the results and are stated here for completeness. First, protein coordinates were made whole across periodic boundaries by a sequential minimum-image walk along the atom ordering before any superposition; without this correction the aqueous trajectory shows a spurious Cα RMSD excursion reaching 9.7 Å that is an artefact of the protein crossing the box boundary rather than a conformational event. Second, in the annular analysis of membrane deformation, bins containing fewer than three phosphorus atoms in a given leaflet were discarded for that frame; the innermost bin (0–12 Å) of the explicit-bilayer system is occupied by the protein itself and never reaches this threshold, which is why it is absent from Fig. 5b. A third detail concerns the bilayer production run. Its 10–20 ns coordinate segment was overwritten on 10 September 2025 by a re-run that was interrupted at 4.18 ns; the GROMACS logs show segments 1, 3, 4 and 5 completing 10 ns each while segment 2 does not. The re-run left five frames covering 10–14 ns, which are discarded because they do not continue the original trajectory; the retained coordinates therefore span 0–10 and 20–50 ns, and the 44 frames they contain are what the analysis uses. Because the equilibrated averages are taken over the second half of the saved frames, they fall entirely after 30 ns and are unaffected; the trace in Fig. 4 is drawn with a break across the missing interval rather than interpolated.

The reverse-QTY substitution set is fully determined by the DSSP assignment of the starting model and can be regenerated from the native sequence without additional information: convert every Q, T and Y whose DSSP code is H, G or I to L, V and F respectively, and leave all others unchanged.
